# A double negative post-transcriptional regulatory circuit underlies the virgin behavioral state

**DOI:** 10.1101/2020.12.16.423061

**Authors:** Daniel L. Garaulet, Albertomaria Moro, Eric C. Lai

## Abstract

The survival and reproductive success of animals depends on the ability to harmonize their external behaviors with their internal states. For example, females conduct numerous social programs that are distinctive to virgins, compared to post-mated and/or pregnant individuals. In *Drosophila*, the fact that this post-mating switch is initiated by seminal factors implies that the default state is virgin. However, we recently showed that loss of miR-iab-4/8-mediated repression of the transcription factor Homothorax (Hth) within the abdominal ventral nerve cord (VNC) causes virgin females to execute mated behaviors. To elucidate new components of this post-transcriptional regulatory circuit, we used genomic analysis of *mir-iab-4/8* deletion and *hth*-miRNA binding site mutants (*hth[BSmut]*) to elucidate *doublesex* (*dsx*) as a critical downstream factor. While Dsx has mostly been studied during sex-specific differentiation, its activities in neurons are little known. We find that accumulation of Dsx in the CNS is highly complementary to Hth, and downregulated in miRNA/*hth[BSmut]* mutants. Moreover, virgin behavior is highly dose-sensitive to developmental *dsx* function. Strikingly, depletion of Dsx in SAG-1 cells, a highly restricted set of abdominal neurons, abrogates female virgin conducts in favor of mated behavioral programs. Thus, a double negative post-transcriptional pathway in the VNC (miR-iab-4/8 -| Hth -| Dsx) specifies the virgin behavioral state.

## Introduction

Females of diverse invertebrate and vertebrate species coordinate multiple behavioral programs with their reproductive state. Mature female virgins are more receptive to male courtship and copulation, but following mating and/or pregnancy, decrease their sexual activity and modulate behaviors to generate and foster their children. Behavioral remodeling associated with the female reproductive state includes increased aggression and nest building in avians and mammals (Ogawa and Makino, 1984; Svare et al., 1982), and decreased male acceptance, increased egg-laying, and manifold appetitive and metabolic changes in insects (Anholt et al., 2020). The genetic and neurological control of this process has been intensively studied in female fruitflies, where sexual activity induces the post-mating switch, a host of behavioral changes collectively known as post-mating responses (PMRs) (Anholt et al., 2020). While some alterations of female behaviors are required for effective reproduction and/or overtly benefit progeny, her reproductive behaviors can also be modulated by the male in ways that are antagonistic to her interests (Anholt et al., 2020). The existence of sexual conflicts within this regulatory program highlights its non-intuitive operation and evolution. Overall, the molecular and circuit mechanisms that control female virgin and mated behaviors are still being elucidated.

In *Drosophila* as in many other species, “virgin” is typically considered the default behavioral state, since factors that induce the post-mating switch are transferred within seminal fluids during copulation. Amongst these, Sex Peptide (SP) is necessary and sufficient to drive most female post-mated behaviors (Kubli and Bopp, 2012). SP signals via uterine SP sensory neurons (SPSNs), which project to the abdominal region of the ventral nerve cord (VNC) (Feng et al., 2014; Hasemeyer et al., 2009; Yang et al., 2009). Some SPSN+ neurons contact abdominal interneurons that express myoinhibitory peptide (Jang et al., 2017), which input into a restricted population of ascending neurons (SAG) that project to the posterior brain including pC1 neurons (Feng et al., 2014; Soller et al., 2006; Wang et al., 2020b). This outlines an ascending flow of information for how a sperm peptide can alter brain activity. Then, the brain integrates this along with auditory and visual cues and coordinates an array of behavioral responses mediated by distinct lineages of descending neurons, and VNC populations that promote or inhibit specific behaviors according to the internal state and the external stimuli (Mezzera et al., 2020; Wang et al., 2020a; Wang et al., 2020b; Wang et al., 2021).

Recently, we made surprising genetic observations regarding implementation of the female virgin behavioral state. In particular, post-transcriptional suppression of the homeobox gene *homothorax* (*hth*) within the VNC is critical to implement the virgin behavioral state (Garaulet et al., 2020). Of note, deletion of the Bithorax Complex (BX-C) miRNA locus (*mir-iab-4/8*), point mutations of BX-C miRNA binding sites in *hth*, or deletion of the *hth* neural-specific 3’ UTR extension bearing many of these miRNA sites, all cause mutant female virgins to perform a host of behaviors characteristic of subjectively mated flies. While mutants of SP and SPR cause mated females to continue to execute virgin behaviors, these represent the first mutants that exhibit the reciprocal phenotype. Thus, the failure to integrate two post-transcriptional regulatory inputs at a single target gene prevents females from appropriately integrating their sexual internal state with external behaviors.

Our recognition of the transcription factor Hth as a target of regulatory circuits for virgin behavior implies the existence of downstream loci that serve as a functional output for this process. In this study, we used molecular genetic profiling to identify a critical requirement for Doublesex (Dsx) to implement the female virgin behavioral state. Although Dsx has been well-studied with respect to the differentiation of sexually dimorphic traits (Kopp, 2012), its roles in post-mitotic neurons are little known. Here, we show that expression of Dsx in the VNC is required for appropriate behavior of virgin females, and that modulation of Dsx in only a few abdominal VNC neurons is sufficient to convert the suite of female virgin behaviors into mated conducts.

## Results

### VNC-iab-8 domain transcriptomes of BX-C miRNA and *hth*-miRNA binding site mutants

Our findings of a similarly defective post-mating switch in deletions of BX-C miRNAs and of post-transcriptional mutants of *hth* beg the question of their downstream functional consequences. We sought insights using transcriptome profiling of the abdominal VNC. The bidirectionally transcribed BX-C miRNA locus encodes two distinct miRNAs: *mir-iab-4* and *mir-iab-8*, that are expressed in Hox-like patterns along the anterior-posterior axis during fruitfly embryogenesis (**Supplementary Figure 1**) (Bender, 2008). GFP sensors reveal that both miRNAs are active in adjacent domains of the abdominal VNC from embryo to adult, with miR-iab-8 deployed in segments posterior to A7 (Bender, 2008; Garaulet et al., 2014; Gummalla et al., 2012; Tyler et al., 2008) (**Figure 1A**, referred hereafter as the iab-8 domain). Expression of the BX-C Hox gene abd-A largely overlaps miR-iab-4 and demarcates the miR-iab-8-5p activity domain (**Figure 1B**). In flies deleted for *mir-iab-4/8* (trans-heterozygous *Δ/C11* mutants), ectopic Hth proteins accumulate in both miR-iab-4 and miR-iab-8 domains from larval stages (**Figure 1C**) to adult (Garaulet et al., 2014; Garaulet et al., 2020). Specific mutations of all miR-iab-4/8 binding sites in the 3’ UTR of the homeodomain-encoding isoform of *homothorax* (*hth[BSmut]*) also derepress Hth protein (**Figure 1C**). Although genetic evidence indicates both miRNAs contribute to the *Δmir-iab-4/8* phenotype in female virgins, ectopic Hth in *hth[BSmut]* is most overt within the miR-iab-8 domain (**Figure 1C** and **H**), and is sufficient to induce PMRs in virgin females (Garaulet et al., 2020).

**Figure 1.**
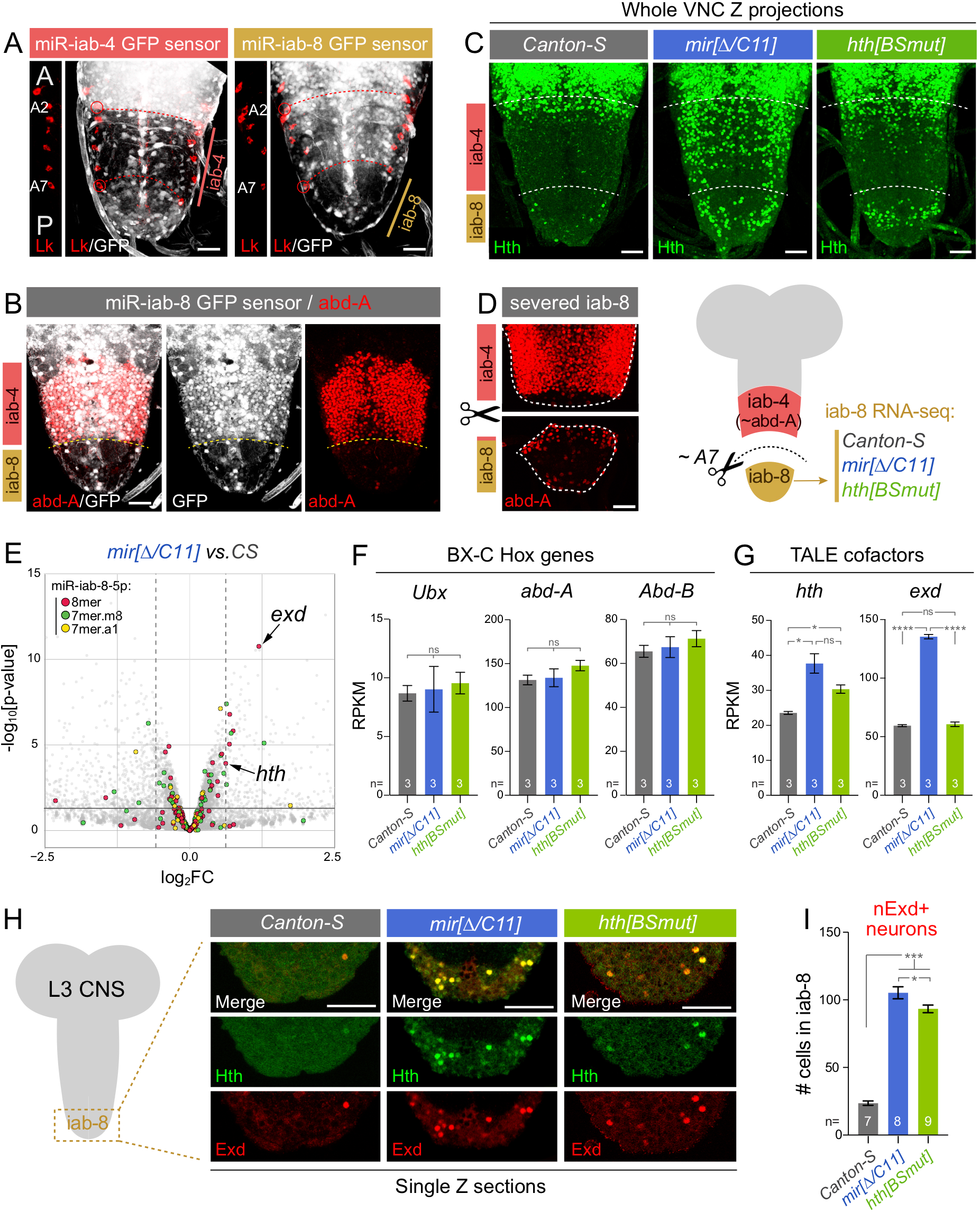
Cytological and transcriptome analysis of a key miRNA/target regulon in the VNC. (A) The bidirectionally transcribed Bithorax Complex (BX-C) miRNA locus yields *mir-iab-4* and *mir-iab-8* from opposite strands. Their activity is associated with repression of ubiquitously transcribed tub-GFP-miRNA sensors, revealing distinct domains in abdominal segments, whose registers are marked by segmental marker Leucokinin (Lk, red). This places miR-iab-4-5p activity in segments A2-A7 and miR-iab-8-5p in A8-A9. A portion of the Lk staining is shown to the left of each image to reference the segment locations; A=anterior, P=posterior. (B) The posterior limit of abd-A (A7a/p) corresponds to the miR-iab-4/miR-iab-8 border. (C) Transcription factor Homothorax (Hth) is a key BX-C miRNA target. It is largely absent throughout the abdominal VNC in wild-type flies, and is derepressed in deletion of BX-C miRNA locus or an engineered allele of *hth-HD* with point mutations in all miR-iab-4/8 binding sites (*hth[BSmut]*). Derepression of Hth protein in the latter is most overt within the iab-8 domain. (D) Validation of dissection strategy to prospectively isolate A8-A9 VNC domain (iab-8 region) for transcriptomics. These stainings are from an individual dissected VNC, but were imaged separately for this montage. (E) Volcano plot of transcripts >1RPKM showing that TALE cofactors *hth* and *exd* are amongst the most significantly derepressed miR-iab-8-5p targets in the BX-C miRNA mutant VNC; most other direct targets were unaffected. The black horizontal line demarcates the 0.05 cutoff for p-value, dotted vertical lines indicate the 1.5 cutoff for fold change. (F-G) Comparison of BX-C Hox genes and TALE cofactors expression in wild-type, miRNA mutant and *hth[BSmut]* iab-8 domain VNCs. *exd* is derepressed in the miRNA mutant but not in *hth[BSmut]*. (H) Single plane confocal images demonstrating nuclear colocalization of Hth and Exd in the iab-8 domain of *mir-iab-4/8* and *hth[BSmut]* VNC. (I) Quantification of ectopic nuclear Exd cells in the iab-8 domain of VNCs of the indicated genotypes. Statistical significance was evaluated using unpaired t test with Welch’s correction. ns, not significant, *p < 0.05, *** p < 0.001, ****p < 0.0001. Error bars, SEM. Scale bars A-D, 25 μm. Scale bars in H, 40 μm.

Based on this, we sought to collect RNA-seq data from the iab-8 domain of females of control VNC and the two mutants. As we lack markers that permit positive selection of this region, we opted for manual separation. We chose to analyze the larval VNC, which owing to its more extended morphology than its adult counterpart, was more amenable to microdissection. With practice, we could reproducibly sever the VNC at the level of A7 pair of nerves, posterior to the major domain of abd-A, as assessed by *post hoc* immunostaining (**Figure 1D**), thus liberating the iab-8 region of the VNC (segments A8 and A9). We prepared triplicate RNA-seq samples from this region from the three genotypes (**Figure 1D** and **Supplementary Figure 1**).

While both mutants were reproducibly distinct from *Canton-S* control, they exhibited limited overall changes in gene expression (**Supplementary Figure 1**). We examined the *Δmir-iab-4/8* data with respect to different classes of conserved seed matches for miR-iab-8-5p (www.targetscan.org). Amongst genes expressed at a minimum level (>1RPKM), the strong majority bearing target sites were unchanged (**Figure 1E** and **Supplementary Figure 1**), and only modestly more targets were upregulated than downregulated (at 1.5-fold change, 10 up and 6 down, **Figure 1E**). Bulk tissue sequencing might underestimate target responses if they were heterogeneous on a cell-by-cell basis. For example, we did not observe significant changes in validated Bithorax-Complex Hox gene targets *Ubx* and *abd-A* (Tyler et al., 2008) (**Figure 1F**), which detectably express ectopic, although sporadic, proteins within the iab-8 domain of BX-C miRNA mutants (Bender, 2008; Garaulet et al., 2014; Gummalla et al., 2012). In any case, as the effects of BX-C miRNA deletion on its targets in the abdominal VNC were limited, it was notable that two of the highest and most significantly upregulated miR-iab-8-5p targets were *hth* and *extradenticle (exd)* (Garaulet et al., 2014) (**Figure 1E, G**). Consistent with detection of ectopic Hth protein, the iab-8 region of *hth[BSmut]* VNC also derepressed *hth* but did not upregulate *exd* transcripts, whose levels were only changed in *Δmir-iab-4/8* (**Figure 1G** and **Supplementary Figure 1**).

Hth and Exd are heterodimeric TALE class homeodomain factors that act as Hox gene cofactors, but also have independent functions. Hth is a spatially patterned nuclear factor, and in the VNC, the anterior boundary of iab-4 expression is normally coincident with the loss of Hth (Figure 1C) (Garaulet et al., 2020). Exd is expressed more broadly, but remains cytoplasmic in the absence of Hth, which serves as its nuclear escort (Pai et al., 1998; Rieckhof et al., 1997). Because of this, nuclear Exd is very sparse in wild-type abdominal VNC segments. In contrast, BX-C miRNA mutants broadly exhibit ectopic nuclear Exd within both iab-4 and iab-8 domains (**Figure 1H**) (Garaulet et al., 2014). If this required joint release of both genes from miRNA control, we might expect a different pattern of abdominal Exd in *hth[BSmut]*. However, the nuclear intensity of ectopic Exd colocalized to that of Hth, and was similar between the two mutant genotypes, despite the fact that *exd* RNA levels were only increased in *mir[Δ/C11]* but not in *hth[BSmut]* mutant VNCs (**Figure 1G** and **1H**). We quantified the iab-8 domains of the three genotypes of overtly Exd+ nuclei, and observed comparable, strong increases in both *mir[Δ/C11]* and *hth[BSmut]* mutants (**Figure 1I**). This suggests that even though Exd is a prominent miR-iab-4/8 target (**Figure 1E**) (Garaulet et al., 2014), it is not limiting for the capacity of derepressed Hth to exert phenotypic or regulatory effects in these mutants.

Given the restricted effects of BX-C miRNA loss on direct targets in the iab-8 VNC, we examined features of presumably indirect changes in gene expression. If derepressed Hth was responsible for some of these effects, they might be associated with overlapping responses between miRNA deletion and *hth[BSmut]* VNC. Intriguingly, we observe substantial overlaps between their up- and down-regulated gene sets, with relatively few genes exhibiting discordant behavior between these two mutants (**Figure 2A** and **Supplementary Table 1**). Compelling loci that are co-regulated between these mutants include neuronal receptors, channels and peptide hormones (**Supplementary Figure 2**). Taken together, these data were potentially consistent with the possibility that deregulated Hth may in large part be responsible for driving aberrant gene expression downstream of loss of BX-C miRNAs.

**Figure 2.**
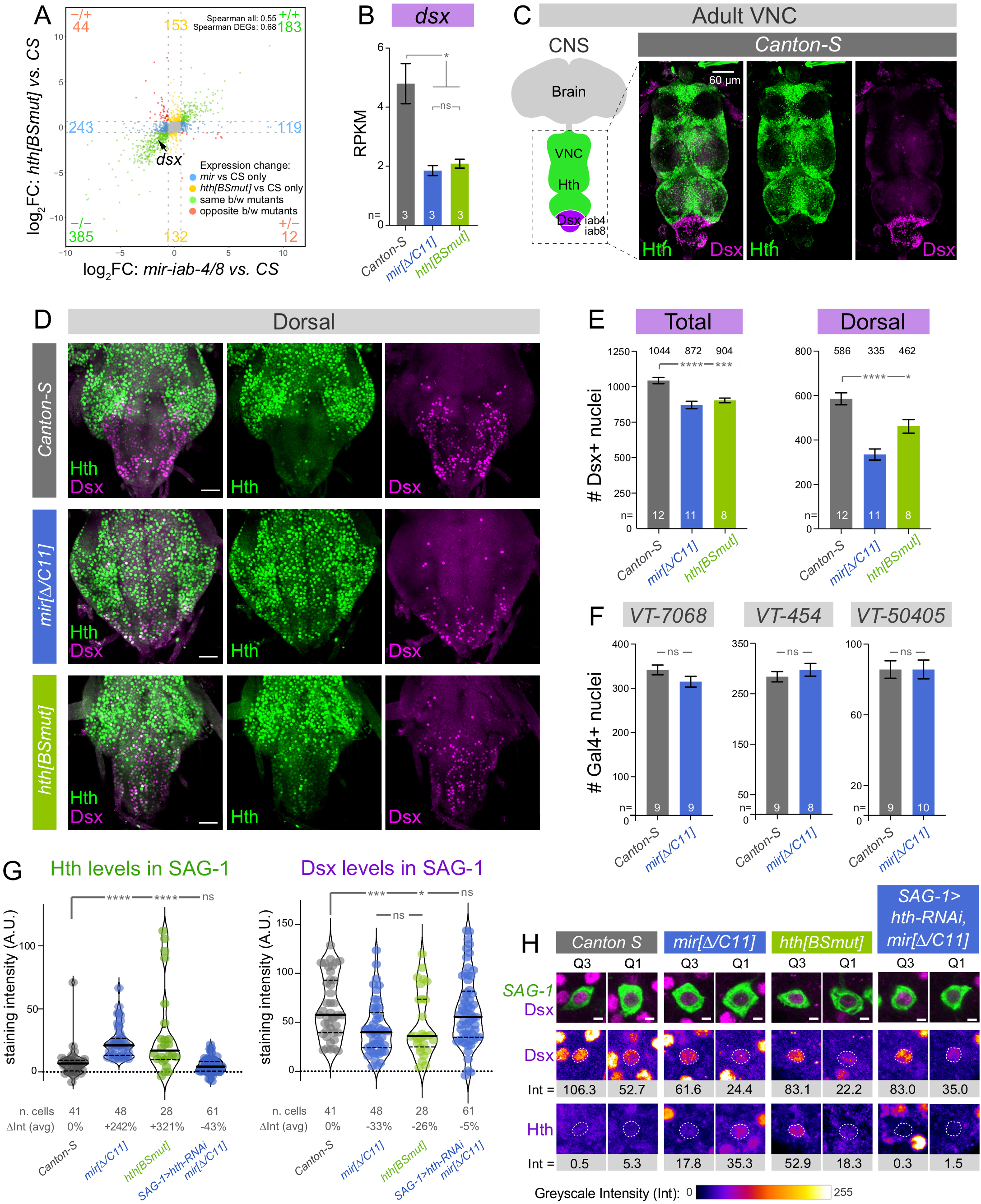
A double negative post-transcriptional circuit: BX-C miRNAs prevent Hth from excluding Dsx in the VNC. (A) Transcriptome analysis reveals substantial similarity in the iab-8 domain of the BX-C miRNA and *hth[BSmut]* mutants, compared to *Canton-S*. Only genes >1RPKM and p<0.05 are plotted. Dotted grey lines indicate fold change = 1.5. (B) A notable downregulated transcript in both mutants is the sex-specific differentiation factor *doublesex* (*dsx*). (C) Hth and Dsx proteins are spatially complementary in the VNC, with Dsx expressed in abdominal segments where the BX-C miRNAs are present. (D) Derepression of Hth in BX-C miRNA mutant (*mir[Δ/C11*]) and *hth[BSmut]* compromises accumulation of Dsx in abdominal segments, especially in dorsal planes as shown here. (E) Quantification of Dsx+ nuclei in wild-type, BX-C miRNA mutant and *hth[BSmut]* VNC; total VNC and dorsal half are shown. The number Dsx+ neurons is compromised in the mutant conditions. (F) Conversely, the number of other identified abdominal subpopulations that mediate PMRs (VT-switch lines) are not affected by miRNA deletion. (G) Quantification of Hth and Dsx protein levels in individual abdominal neurons relevant to the female post-mating switch (abdominal SAG-1 neurons). Depression of Hth correlates with reduction of Dsx within identified neurons. (H) Representative GFP-labeled SAG-1 neurons from higher (Q1) and lower (Q3) quartiles of Dsx, as quantified in (G), with corresponding Dsx and Hth levels. The nuclear levels of each factor (Int) were calculated by subtracting the signals in cytoplasm (marked in green) from the corresponding nucleus (dotted line) for each antigen; samples were co-stained and imaged in parallel within the linear range. (B,E,F) t-test with Welch’s correction; (G) Mann-Whitney non-parametric test. ns, not significant, *p < 0.05, ***p < 0.001, ****p < 0.0001. Error bars, SEM. Scale bars, 60 μm in C, 30 μm in D, 2 μm in H.

### Spatial complementarity of Hth and Dsx is disrupted by loss of miRNA regulation

Amongst the genes co-regulated by BX-C miRNA loss and deletion of their binding sites from *hth*, we were particularly intrigued by *doublesex* (*dsx*), which was ∼2-fold downregulated in both mutants (**Figure 2A-B**). This transcription factor is widely studied for its central role in sex determination locus, and its vertebrate homolog DMRT1 similarly controls sex-specific differentiation (Kopp, 2012). However, less is known about its functions in post-mitotic neurons. We were intrigued by the highly spatially complementary pattern of Dsx and Hth proteins in the nervous system. In the VNC, Dsx protein is restricted to the posterior abdominal ganglion and abuts the domain of Hth, located more anteriorly (**Figure 2C**). The reciprocal pattern of Dsx and Hth also extends to the brain. While Dsx accumulates more sparsely in this setting, only rarely is it Dsx colocalized with Hth, even amongst closely apposed cells (**Supplementary Figure 3**). In the VNC, only very few neurons co-express these proteins (**Supplementary Figure 3**).

Immunostaining of BX-C miRNA mutants revealed a decrease in Dsx+ neurons in the abdominal ganglion of both miRNA and *hth[BSmut]* mutants. This was most visually evident in the dorsal region of the VNC (**Figure 2D**). To be certain of this effect, we quantified all Dsx+ neurons throughout the volume of the VNC, and observed ∼150 fewer Dsx+ neurons in both mutant conditions (**Figure 2E**), but found that the difference was larger when restricting the analysis to the dorsal half of the VNC (**Figure 2E**). We recently showed that repression of *hth* by BX-C miRNAs that is relevant for female PMRs occurs in several populations of abdominal VNC neurons, marked by the *VT-7068, VT-454* and *VT-50405* Gal4 drivers (i.e., VT-switch lines) (Feng et al., 2014; Garaulet et al., 2020). However, the numbers of neurons labeled by these drivers (∼80-300) were not substantially affected in miRNA mutants (**Figure 2F**). Thus, loss of Dsx reactivity in mutants was not due to loss of abdominal neurons that are involved in the post-mating switch *per se*.

The regulatory intersection of *VT-7068* and *VT-50405* (as a split-Gal4 combination) labels a handful of neurons in the entire CNS of female flies, with typically four “SAG-1” abdominal neurons found in the iab-8 domain (**Supplementary Figure 4**) (Feng et al., 2014). These project to the central brain (**Supplementary Figure 4**) and constitute a minimal set of VNC cells whose enforced activation can partially induce virgin behaviors in mated females (Feng et al., 2014). We employed these abdominal SAG-1 neurons for quantitative analysis of Hth and Dsx, taking great care to stain wild-type and mutant VNC in the same wells and to image them in parallel using identical settings (see Methods). Although abdominal SAG-1 neurons did not necessarily accumulate the highest levels of Hth seen amongst total *VT-7068* or *VT-50405* neurons (Garaulet et al., 2020), we could reliably observe elevated Hth in identified SAG-1 neurons, in both miRNA deletion and *hth[BSmut]* mutants, concomitant with downregulation of Dsx (**Figure 2G and 2H**). In these same SAG-1 neurons, wild-type Dsx exhibits a somewhat bimodal distribution (**Figure 2G and 2H**). However, deletion of BX-C miRNAs eliminates most of the Dsx-high class, and total levels of Dsx are significantly lower in both *mir-iab-4/8* and *hth[BSmut]* mutants (**Figure 2G**). To illustrate the diversity, but overall reduction of Dsx levels, we show two representative neurons per genotype, corresponding to the first and third quartiles (Q1, Q3) from **Figure 2H**. The comparison of Dsx intensity (Int) between neurons of the same quartile across wild-type, *mir-iab-4/8* and *hth[BSmut]* mutants reveals lower Dsx accumulation (**Figure 2H**). Therefore, loss of miRNA-mediated regulation of Hth results in its derepression in abdominal VNC neurons, which in turn is associated with decreased Dsx.

To understand if the decrease in abdominal Dsx levels was promoted by elevated Hth, we used the SAG-1 driver to ectopically express *hth* and observe Dsx levels in abdominal SAG-1 nuclei. These flies showed a slight reduction of Dsx accumulation, although the difference with wild-type was not significant (**Supplementary Figure 4**). Conversely, ectopic expression of Dsx protein decreased Hth moderately (**Supplementary Figure 4**). These gain-of-function experiments are difficult to interpret since the levels of overexpressed Hth or Dsx are very high, and likely non-physiological (**Supplementary Figure 4**). Therefore, we utilized a different approach to deplete *hth* in *mir-iab-4/8* mutants. In *SAG-1>hth-RNAi; mir[Δ/C11]* virgins, Dsx levels and distribution were restored to wild-type values (**Figure 2G** and **2H**). This result confirms that the decrease of Dsx protein observed in mutants requires elevation of endogenous Hth. In other words, there is a double negative relationship extending from the miRNA to Hth to Dsx.

### Doublesex is highly dose sensitive for female virgin behavior

*doublesex* is necessary for male courtship (McRobert and Tompkins, 1985) as well as to specify the female circuitry necessary for the post-mating switch (Rezaval et al., 2012; Rideout et al., 2010; Robinett et al., 2010). However, despite the fact that *dsx* expressing neurons are required for the post-mating switch and reproductive behaviors in female flies, the functional impact of Dsx protein in female behavior has not yet been tested.

We initially examined this using independent loss of function alleles of *dsx. doublesex* null mutants display inter-sex cuticular features, including male and female elements on their genitalia (Nothiger et al., 2009). However, *dsx* heterozygotes lack overt cuticular defects. We decided to test female heterozygotes in order to understand the effects of partial *dsx* decrease, in order to mimic the phenotype observed in *mir-iab-4/8* and *hth[BSmut]* mutants. As expected, a single dose of *dsx* gene is sufficient for flies to develop normal genitalia, as well as a normal number and appearance of sex-combs on male forelegs (**Supplementary Figure 5**). These sexually dimorphic structures on male forelegs typically disappear in *dsx* homozygotes in favor of thinner and shorter bristles, characteristic of female flies (**Supplementary Figure 5**). Thus, *doublesex* is haplo-sufficient for gross anatomy.

We then examined virgin behavior. The reception of male sperm during copulation induces profound physiological and behavioral changes in female flies (Anholt et al., 2020). The male component Sex Peptide (SP) triggers a cascade of neuronal events including the inactivation of the minimal core of ascending information to the brain: a few dozen of uterine sensory neurons (SPSN’s) detect the presence of the male seminal protein in the female uterus and transmit this information to the abdominal ganglion (AbG) in the VNC (Feng et al., 2014; Hasemeyer et al., 2009; Yang et al., 2009), where they input to Mip and SAG-1 neurons (Jang et al., 2017). In response to this peripheral input, SAG-1 neurons attenuate their activity, which signals the brain of the mating status of the female reproductive tract (Jang et al., 2017; Wang et al., 2020b). From that point, the central brain integrates this input with visual and auditory cues and coordinates a large remodeling of female reproductive behaviors. As a result, female virgins and inseminated females display largely different suites of behavioral adaptations modulated by distinct descending neuronal lineages (Mezzera et al., 2020; Wang et al., 2020a; Wang et al., 2020b; Wang et al., 2021).

The two most commonly monitored aspects of this behavioral switch are egg-laying and receptivity. Female virgins are highly receptive to male courtship and tend to lay fewer eggs in the first few days after eclosion (**Figure 3A,B**). Following copulation, females remain refractory to further copulation attempts for a few days and increase egg-laying substantially (Anholt et al., 2020) (**Figure 3A,B**). Surprisingly, *dsx* heterozygosity compromises both behaviors: egg-laying is increased and receptivity is decreased compared to wild-type virgins. Importantly, both of these readouts were similarly and significantly affected in independent dsx alleles, compared to their control siblings (**Figure 3A,B**). Together, these slight but genetically robust differences in egg-laying and receptivity seemed to indicate a partial transition to the mated state in pre-inseminated virgins triggered by *dsx* heterozygosity. To confirm this hypothesis, we tested two additional behaviors that are associated with female internal states. Vaginal plates opening are mostly performed by receptive virgins, and very rarely observed in the early days after copulation (Wang et al., 2021) (**Figure 3C**). Conversely, mating elicits a specific form of rejection of males consisting in the maintained, full extrusion of the ovipositor when a male is actively courting (Mezzera et al., 2020; Wang et al., 2020a) (**Figure 3D**). For both of these performances, independent *dsx* heterozygotes also differ from canonical wild-type virgins. Together, *dsx* heterozygosity attenuates virgin behaviors while simultaneously enhancing mated-specific PMRs, suggesting the subjective induction of the post-mated state (**Figure 3A-D**).

**Figure 3.**
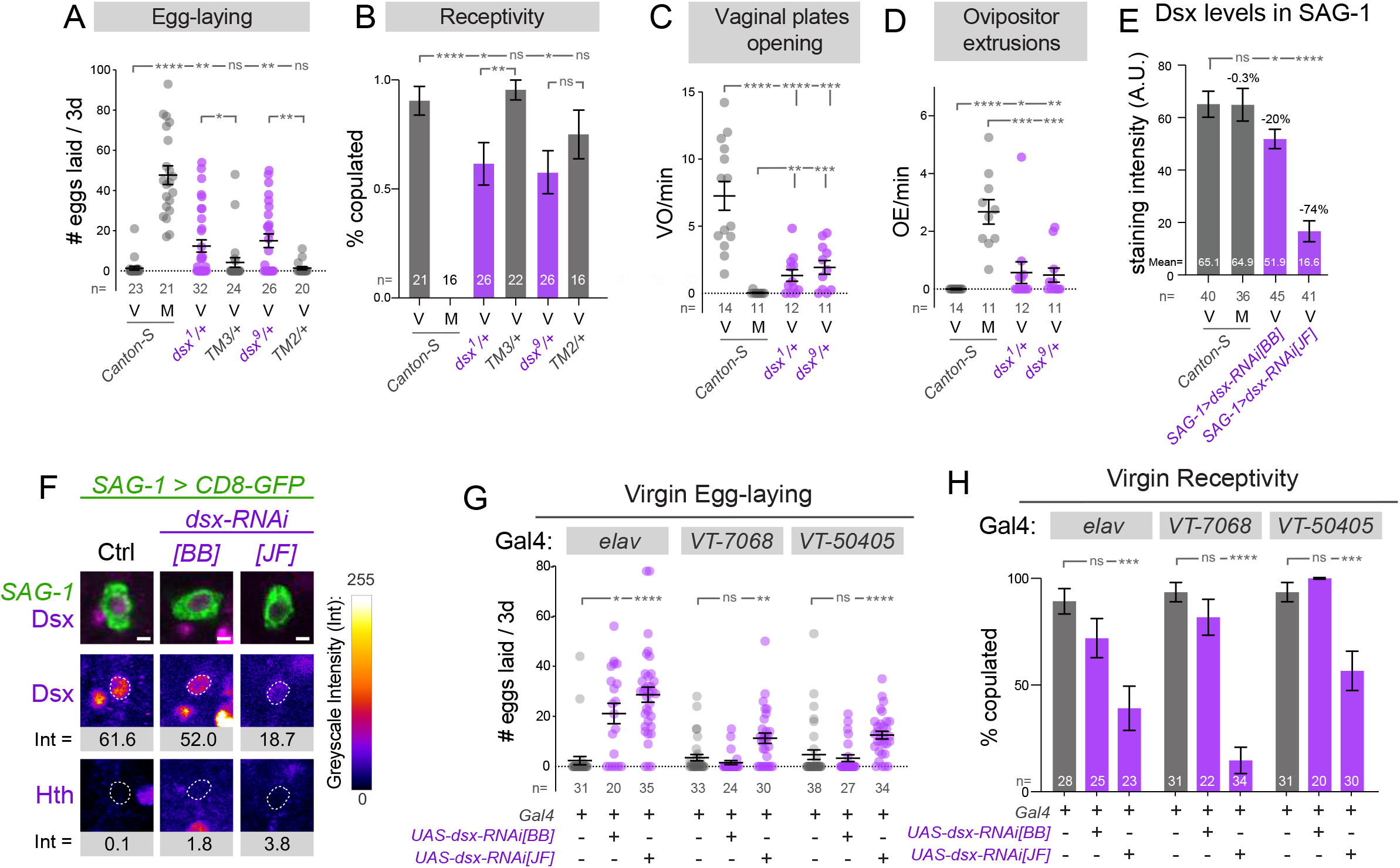
Endogenous Dsx is essential for females to interpret the virgin behavioral state. (A-D) Comparison of female behaviors in wild-type virgins, mated females, and *doublesex* heterozygotes, illustrating a partial transition to the mated state in the latter. (E) Analysis of Dsx levels in abdominal SAG-1 neurons pre- and 24h post-insemination. Validation of independent *dsx-RNAi* transgenes. *dsx-RNAi[JF]* has a stronger effect on Dsx in SAG-1 neurons. (F) Representative GFP-labeled abdominal SAG-1 neurons from average Dsx levels are shown, with corresponding Dsx and Hth staining. The dotted line corresponds to the nucleus of each neuron. (G, H) Pan-neuronal (using *elav-Gal4*) knockdown of *dsx* using *[BB]* and *[JF] UAS-dsx-RNAi* lines results in increased egg-laying and decreased receptivity by female virgins. Expression of the *dsx-RNAi[JF]* transgene in restricted sets of abdominal VT-switch neurons also induces virgin egg-laying and reduces receptivity. (A,C,D,E,G) Mann-Whitney non-parametric test; (B,H) Fisher’s exact test. ns, not significant, *p < 0.05, **p < 0.01, *** p < 0.001, ****p < 0.0001. Error bars, SEM. Scalebar in F, 2 μm. Eggs were collected over 3 days for all virgin genotypes and over the first 24h after copulation in mated flies. V=virgin females, M=mated females.

Given that female virgin performance depends on *dsx* dosage, we then wondered if PMRs in wild-type females could be triggered by a *dsx* decrease as a putative effect of mating. We analyzed Dsx accumulation in abdominal *SAG-1* neurons, the key element of the ascending post-mated switch circuit (Feng et al., 2014). By applying standardized staining and imaging of female VNC, we could not find any difference in Dsx intensity within the two states: virgins and females after 24 hours from insemination. Although we cannot rule out that other neuronal lineages might be experiencing a fluctuation in Dsx levels, our results suggest that Dsx is not an active component of the switch in wild-type flies. Similarly, we previously showed that Hth levels in SAG-1 neurons do not change either in response to mating but instead, their regulation is crucial for abdominal neurons during development (Garaulet et al., 2020). Our results here are compatible with the hypothesis that *dsx* functions during development might be critical to imprint the right anatomy or function of switch neurons that will be required in adults, rather than an active regulation of their function in response to mating. In fact, although other genes like *fruitless* and *pickpocket* are expressed in some of the neurons regulating female reproductive behaviors, Dsx is found in most of the lineages comprising the ascending circuit (SPSNs, Mip, SAG-1) (Feng et al., 2014; Jang et al., 2017) as well as the descending circuit controlling egg-laying, ovipositor extrusions and vaginal plates openings (pC1, pC21, DNp13, OviDNs) (Mezzera et al., 2020; Wang et al., 2020a; Wang et al., 2021).

### Expression of Doublesex in restricted abdominal neurons is required for virgin behavior

With knowledge that virgin behavior is highly sensitive to *dsx* dosage, we next interrogated where and when Dsx is required in the nervous system for female responses. For this purpose, we used a stock published by Bruce Baker’s lab (*dsx-RNAi[BB]*) (Robinett et al., 2010) and an independent line from the TRiP-JF collection (*dsx-RNAi[JF]*). Although the *dsx-RNAi[BB]* stock actually contains two copies of this transgene, it was documented to induce only weak *dsx* knockdown at 25° C (Robinett et al., 2010); *dsx-RNAi[JF]* has not been directly assessed for Dsx suppression. We compared their efficacies when activated at 25° C using *SAG-1-Gal4*, which allowed us to carefully quantify Dsx alteration in the defined set of abdominal SAG-1 cells where *dsx* is normally expressed. We found that *dsx-RNAi[BB]* induced mild but significant reduction of Dsx protein, within the range observed in the miRNA and *hth[BSmut]* mutants, while *dsx-RNAi[JF]* induced stronger depletion (**Figure 3E, F** compare to **2G, H**). Examples of representative stained cells are shown in **Figure 3F**. Consistent with this hierarchy, *dsx-RNAi[JF]* but not *dsx-RNAi[BB]* could reduce the number of male sex comb bristles at 25°C (**Supplementary Figure 5**). Of note, despite its ability to induce substantial knockdown, *dsx-RNAi[JF]* induced only a mild cuticular defect. By comparison, the viable trans-allelic combination of *dsx[1]/[9]* results in full loss of differentiated male sex combs (**Supplementary Figure 5**), as well as general inter-sex features. Thus, we have inducible genetic reagents that reproduce mild suppression of Dsx protein and function, within a range that is relevant to disruption of Dsx levels that occurs in BX-C miRNA deletion and *hth[BSmut]* alleles.

We then used egg-laying and receptivity as readouts to screen a potential shift to a subjective post-mated state. Upon induction of either *dsx-RNAi* transgene with pan-neuronal *elav-Gal4*, we were excited to observe that both *dsx-RNAi* transgenes reliably increased the egg-laying capacity of young female virgins (**Figure 3G**), in line with the results obtained for *dsx* heterozygotes. Similarly, receptivity in these two genotypes was significantly compromised (**Figure 3H**). We additionally tested fertility and climbing in these genetic combinations to rule out general, non-specific effects of *dsx* depletion, an undesired side effect of RNAi machinery activation, or possibly off-targeting. Both male and female fertility and locomotor activity in the mutant combinations were indistinguishable from controls, emphasizing the specificity of *dsx* depletion on virgin behaviors (**Supplementary Figure 5**). Of note, our results with *dsx* heterozygotes and knockdowns highlight the sensitivity of behavior to *dsx* dose, in comparison to cuticular structures, which remain unperturbed under similar *dsx* depletion.

Restriction of *dsx* knockdown to the VT-switch lineages using *VT-7068* and *VT-50405* Gal4 lines induced similar effects on both egg-laying and receptivity (**Figure 3G,H**). Interestingly, in all cases the phenotypes effects observed with the stronger line *dsx-RNAi[JF]* were stronger than with *dsx-RNAi[BB]*, further highlighting *dsx* dosage sensitivity of virgin behaviors. Then, we tested if modulation of Dsx in the intersection of these two drivers, the sparser SAG-1 lineage, was sufficient to stimulate egg-laying and suppress virgin receptivity.

Indeed, both RNAi lines induced aberrant egg-laying in young virgins, and at least *dsx-RNAi[JF]* compromised virgin receptivity to values that are comparable to both mutant conditions (**Figure 4A, B**). These phenotypes were in line with those observed in *mir-iab-4/8* and *hth[BSmut]* mutants (Garaulet et al., 2020) (**Figure 4A, B**), suggesting that Dsx decrease could be causal to behavioral defects in the absence of *hth* regulation by miR-iab-4/8. To investigate if depletion of *dsx* generally affected differentiation, we analyzed the dendritic (VNC) and axonal (brain) pattern of SAG-1 ascending neurons in wild-type controls and the two knockdown conditions. We did not observe any gross anatomical defects upon *dsx* depletion (**Supplementary Figure 5**). Together with *dsx* heterozygotes and fertility and climbing controls in *elav-Gal4>UAS-dsx-RNAi*, our experiments indicate that virgin behaviors and PMRs are specifically sensitive to Dsx levels. We note that the split-Gal4 system may confer enhanced activity over the individual VT-switch lines. But, the fact that the knockdown occurs in very restricted neurons, including the four ascending neurons (**Figure 3E**) and is only a partial knockdown, indicates a strong requirement for Dsx in these cells for female behavior.

**Figure 4.**
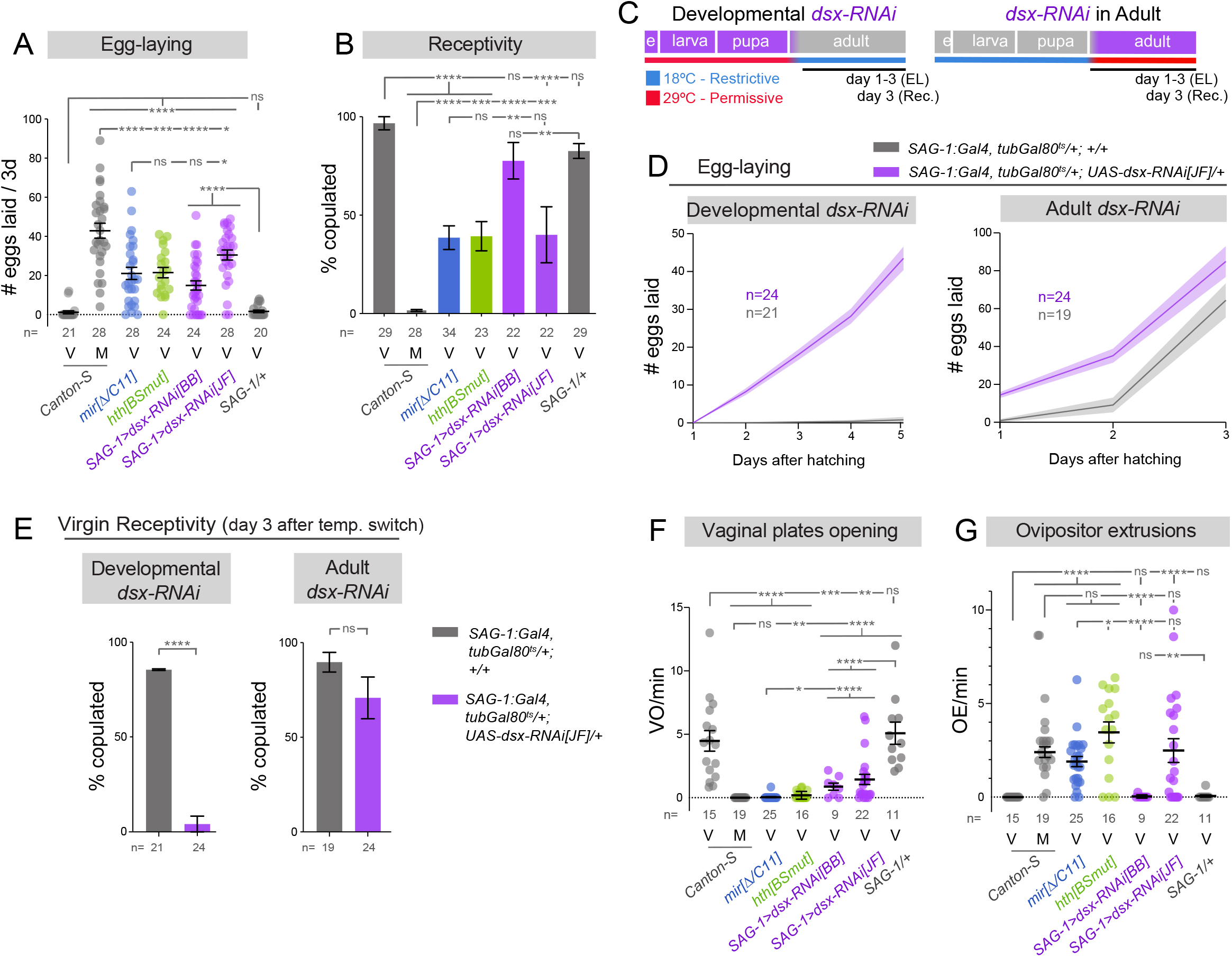
SAG-1 neurons specifically require Dsx for a suite of female virgin behaviors. (A,B) Suppression of Dsx within the SAG-1 lineage enhances egg-laying and compromises virgin receptivity, similar to whole-animal *Δmir-iab-4/8* and *hth[BSmut]* mutants. *dsx-RNAi[JF]* has a stronger effect. (C) Strategy for temporally-restricted depletion of *dsx*. (D-E) Analysis of egg-laying (D) and receptivity (E) in flies under either developmental or adult knockdown of *dsx*, highlighting the developmental role of *dsx* for virgin adult behaviors. (F, G) Modulation of Dsx levels also decreases vaginal plates opening (F) and induces ovipositor extrusions (G), comparable to deletion of BX-C miRNAs and mutation of BX-C miRNA binding sites in *homothorax (hth[BSmut])*. (A,F,G) Mann-Whitney non-parametric test; (B, E) Fisher’s exact test. ns, not significant, *p < 0.05, **p < 0.01, *** p < 0.001, ****p < 0.0001. Error bars, SEM. Eggs were collected over 3 days for all virgin genotypes and over the first 24h after copulation in mated flies. V=virgin females, M=mated females.

Having identified a minimal set of neurons were *dsx* is necessary for egg-laying and receptivity, we sought the timing of Dsx activity to regulate these behaviors. Our RNA-seq experiments demonstrate differential expression of *dsx* in miRNA and Hth[BSmut] mutants in late third instar larvae, prior to the establishment of the adult circuitry controlling PMRs. We temporally restricted *dsx* knockdown by including temperature sensitive *Gal80* (*tubGal80*^*ts*^) transgene in *SAG>dsx-RNAi[JF]* flies, and switching them from the restrictive (18 °C) to permissive (29 °C) temperatures, and vice versa right after at eclosion (**Figure 4C**). These temperature shifts allowed us to discern between developmental or adult functions of Dsx in SAG-1 neurons in relation to virgin behaviors and PMRs. Monitoring egg-laying and receptivity, we found that adult knockdown induced subtle, but statistically significant, differences in these two behaviors. In contrast, the effects of developmental knockdown were dramatic for virgin performance: while parameters were unaffected by this temperature regime in controls (little egg-laying and full receptivity), experimental individuals laid eggs profusely and remained completely refractory to male courtship (**Figure 4D, E**).

Our previous work showed that disruption of miR-iab-4/8 regulation of *hth* in the VNC, both in *mir-iab-4/8* or *hth[BSmut]* mutants, suppresses virgin behaviors and induces a switch to a subjective mated state, monitored by at least four different behaviors (Garaulet et al., 2020). This was further confirmed by the fact that just increasing activity of VT-switch neurons in the VNC was sufficient to revert the phenotype in *mir-iab-4/8* mutants, in a similar way as increased activity of these neurons is sufficient to revert the effects of mating and make inseminated females behave as virgins (Garaulet et al., 2020). We tested if depletion of Dsx could potentially cause a broader switch in the behavioral output of virgin females towards a subjective mated state. To this end, in addition of egg-laying and receptivity we characterized a suite of other behavioral analyses in *SAG-1>dsx-RNAi[BB]* and *[JF]* females and compared them to wild-type virgin and mated females, as well as to *mir-iab-4/8* and *hth[BSmut]* mutant virgins. By analyzing virgin behaviors that are defective in mutants (e.g., opening of vaginal plates) as well as mated behaviors that are ectopically gained by mutant virgins (e.g. ovipositor extrusion), we gain greater confidence that depletion of *dsx* reflects an active switch in behavioral state. Indeed, these analyses demonstrate that *SAG-1>dsx-RNAi* females qualitatively fail to coordinate virgin status with virgin behavioral programs, and instead act as mated animals (**Figure 4A,B** and **F,G**). We emphasize that in all of these readouts, the highly cell-restricted depletion of *dsx* at least qualitatively recapitulates the effects of BX-C miRNA deletion as well as the mutation of miRNA sites in *hth-HD* (Garaulet et al., 2020). For most cases using the *dsx-RNAi[JF]*, the effects observed were also quantitatively comparable to *Δmir* and *hth[BSmut]* mutants. Although these results do not imply that SAG-1 is the only lineage affected these mutants, they do reveal that Dsx function in these specific cells is pertinent for normal behavior. In summary, a double negative regulatory axis within the central nervous system is critical for virgin females to appropriately coordinate their external behaviors with their internal state (**Figure 5**).

**Figure 5.**
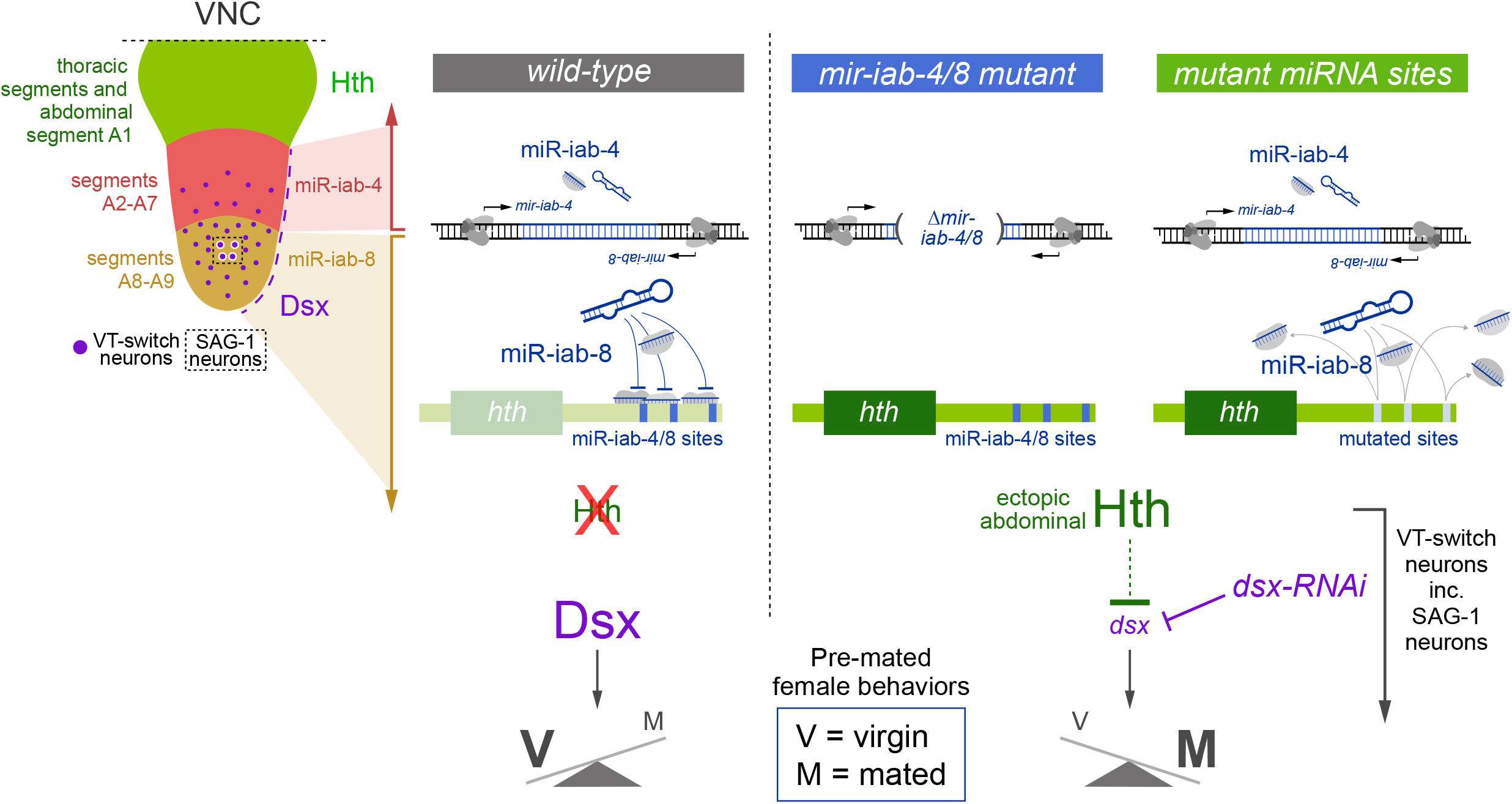
Model for the genetic and spatial control of female virgin behavior in the VNC. Upper left, ventral nerve cord (VNC) illustrates the protein domains of Hth (in thoracic segments and abdominal segment A1) and Dsx (in A2-A9), and RNA domains of *mir-iab-4* (A2-A7) and *mir-iab-8* (A8-A9). Restricted neural populations that govern the female post-mating switch are distributed within A2-A9 (VT-switch neurons), including four SAG-1 neurons that reside within the *mir-iab-8* domain. In the wild-type abdominal VNC, the activity of miR-iab-4/8 suppress Hth, permitting Dsx expression that is critical for execution of virgin behaviors. Virgin females that lack *mir-iab-4/8*, are mutated for these miRNA binding sites within *hth*, or that are depleted of *dsx* within VT-switch neurons (and in as few as those of highly restricted SAG-1 neurons) exhibit a switch from virgin to post-mated behaviors.

## Discussion

Our recent studies established an unexpected requirement for miRNA-mediated suppression of the transcription factor Hth to safeguard the virgin female behavioral state (Garaulet et al., 2020). We used genetically engineered alleles, along with spatial and temporal *hth* manipulations, to demonstrate a developmental requirement for post-transcriptional regulation of Hth within the abdominal ganglion of the CNS for female behavior. However, Hth did not appear to be required in otherwise wild-type VT-switch neurons for execution of virgin behaviors, implying that Hth must be prevented from being expressed in the abdominal VNC. This involves the integration of two regulatory mechanisms, a high density of BX-C miRNA binding sites (miR-iab-4/8) within the *hth-HD* 3’ UTR, as well as a strategy for neural-specific 3’ UTR elongation, which reveals many of these sites only on neural *hth* isoforms.

Here, we extend this novel regulatory axis further, by showing that loss of BX-C miRNAs, acting through derepressed Hth, leads to downregulation of the Dsx in the abdominal VNC. Dsx is well-known as a master sex determination transcription factor (Kopp, 2012), and it shows localized expression in specific CNS domains. However, while the activity of Dsx-expressing neurons *per se* has been implicated in the switch in females (Feng et al., 2014; Hasemeyer et al., 2009; Yang et al., 2009), the functions of Dsx in post-mitotic neurons are less well-defined. Our work reveals that Dsx itself is a new, central component in specifying virgin behavior, since its restricted suppression in as few as four (SAG-1+) neurons is sufficient to induce post-mated behaviors.

Altogether, in contrast to highly branched regulatory networks that are bioinformatically inferred to lie downstream of individual miRNAs, we reveal a double-negative regulatory cascade comprising miRNAs and two transcription factors (**Figure 5**). These findings provide impetus to assess possible direct regulation of Dsx by Hth, as well as to elucidate Dsx targets that are relevant to female behavioral control. Overall, we expand a novel genetic hierarchy that is essential for females to couple the virgin internal state with appropriate behaviors.

## STAR METHODS

### RESOURCE AVAILABILITY

#### Lead contact

Further information and requests for resources and reagents should be directed to and will be fulfilled by the Lead Contact, Eric Lai (, tel: 212-639-5578), or by Daniel L. Garaulet.

#### Materials availability

Transgenic flies generated in this study are available from the corresponding authors on request.

#### Data and code availability

The original behavior data for Figures 1, 3, 4 and Supplementary Figures 5 in the paper are available from the corresponding authors. The raw RNA-seq data reported in this study were deposited in the NCBI Gene Expression Omnibus under accession GSE166562.

## EXPERIMENTAL MODEL AND SUBJECT DETAILS

This study used male and female flies of wild-type and genetically engineered strains of *Drosophila melanogaster*. Virgin and mated parameters refer to assays of female behavioral performances.

### Fly strains and maintenance

Larval and adult flies were raised on cornmeal/molasses media recipe: 83.8% water, 0.6% agar, 4.6% cornmeal, 2.3% dried yeast, 7.8% molasses solids, 0.3% propionic acid, 0.1% tegosept, 0.5% ethanol. They were kept at 25°C (unless mentioned otherwise), 55% humidity and under 12h:12h LD cycles.

*Drosophila* lines used in this study: *mir[Δ]* (Bender, 2008), *mir* [*C11*] and *hth[BSmut]* (Garaulet et al., 2020), *Canton-S* (gift of Karla Kaun), *VT-lines* and *SAG-1* split Gal4 line (Feng et al., 2014), *UAS-hth-RNAi* (Vienna Drosophila RNAi Center), *2xUAS-dsx-RNAi[BB]* (Robinett et al., 2010), *tub-GFP-mir-iab-4 and -iab-8* sensors (Tyler et al., 2008). The following lines were obtained from Bloomington Drosophila Stock Center: *elav-Gal4[C-155]* (BDSC #458), *UAS-mCD8-GFP* (BDSC #5137), *UAS-Red-Stinger* (BDSC #8547), *tubGal80*^*ts*^ (BDSC #7108), *UAS-dsx-RNAi[JF]* (BDSC #26716), *dsx[1]* (BDSC #1679), *dsx[9]* (BDSC #44223), *UAS-dsxF* (BDSC #44223).

All the lines used in this study have been backcrossed at least 8 generations to the *Canton-S* wild-type strain.

## METHOD DETAILS

### RNA extraction and sequencing

Female larvae were dissected on ice for a maximum of 30 minutes (30-40 larvae). The posterior third of larvae was removed with forceps, the remaining was turned inside down. Using 2mm curved blade spring scissors (Fine Science Tools #15000-04), VNC were severed at the level of the A7 pair of nerves, and immediately placed into TRIzol (Thermo Fisher Scientific #15596018). After dissections, samples in TRIzol were stored at -80° C. Each biological replicate pooled severed VNCs of 120-150 larvae, dissected in 4-5 periods of 30 min. 3 replicates were generated per genotype. RNA extraction was performed with TRIzol.

500 ng of total RNA per dissected VNC sample was used for TruSeq stranded mRNA library preparation (Illumina) by the Integrated Genomics Operation (IGO) core at MSKCC. Libraries were sequenced on Illumina HiSeq-1000 sequencer with PE-100 mode. The raw sequence data are available from GEO accession number: GSE166562.

### Bioinformatic analysis

RNA-seq data were aligned to the *Drosophila melanogaster* reference genome (Version r6.21) using HISAT2 software with standard parameters (Kim et al., 2015). We used featureCounts in the Rsubread package to compute features and read numbers for each bam file (Liao et al., 2019). The read counts per gene were then normalized to obtain RPKM values using edgeR package from R Bioconductor (Robinson et al., 2010) and extracting transcript length from Biomart (Durinck et al., 2005). Fold changes between samples were calculated using edgeR applying no filter. Genes targeted by miR-iab-8-5p were identified using conserved TargetScanFly predictions (Agarwal et al., 2018).

### Immunohistochemistry, imaging, and image quantification analysis

Larval and adult CNS were dissected in cold PBS and fixed for 1h in 4% paraformaldehyde + 0.1% Triton. Primary and secondary antibodies were incubated for >36h at 4°C in wash buffer (PBS + 1% BSA) and mounted in Vectashield (Vector Labs). Antibodies used were mouse anti-abd-A (gift of Ian Duncan), rabbit and guinea pig anti-Hth (Salvany et al., 2009), rat anti-Dsx (Sanders and Arbeitman, 2008) and Alexa-488, -555, -647 conjugated goat and/or donkey antibodies from Thermo Fisher Scientific.

Imaging was performed in a Leica TCS SP5 confocal microscope. Each VNC was typically scanned in 55 planes (Z step ∼2 µm). When image quantification or comparison was performed (**Figure 3**), all different genotypes used were dissected at the same time, fixed and incubated together in the same well. To identify the genotype of each VNC while mounting, different parts of the head (eyes, proboscis, antennae, etc.) were left attached or removed from the VNC during dissection. Then, the same number of VNCs from different genotypes were arranged in a known fashion per slide, to avoid differences in the quantification due to the mounting process. Laser power and offset were maintained identically for all the samples being compared. Gain was slightly adjusted to an internal control in each case.

Image quantification analysis was performed using FIJI (Schindelin et al., 2012). To obtain the values of Dsx intensity, each nuclei was identified on the GFP channel, its Dsx/Hth signal measured, and individually normalized by subtracting the background signal of a similar area in the cytoplasmic region of the same cell. All adult VNC images are 0-24 hr old females.

### Behavioral assays

We collected virgin males and females after eclosion and kept them isolated in vials at 25° C, 55% humidity and 12h:12h LD cycles until utilized for behavior assays. All tests were performed at ZT 7-11 and at least at four different occasions. Vaginal plate opening, ovipositor extrusions and receptivity were assayed at day 3 after eclosion in custom 18-multiplex mating arenas (chamber size: 10 mm diameter). From eclosion to day 3, individual male and female virgins were kept isolated in vials. At day 3, single males and females were placed in a half of each arena and allowed to acclimate for 5 min before the assay. Then, they were allowed to interact and recorded for 10 minutes. Ovipositor extrusions and vaginal plates openings were analyzed during the first 4 minutes after courtship initiation or until mating. Counts of either behavior were normalized to time (min). Receptivity was calculated as the cumulative proportion of animals mated at 10 min. Egg-laying was calculated as the number of eggs laid in the first 3 days after eclosion (for virgins of all genotypes), and during the first 24h after copulation (for mated females).

For mated behaviors, virgin females were kept isolated in individual vials. At day 3, they were allowed to mate to CS males, and immediately separated from males after copulation. Then, they were placed individually in single vials. Egg-laying in mated females is the number of eggs laid per female during the first 24h after mating. At 24h after mating, mated females were assayed for vaginal plate opening, ovipositor extrusions and receptivity with fresh males, following the protocol detailed above.

For temperature shifts, flies carrying tub-Gal80-ts were placed at either restrictive (18°C) or permissive (29°C) during development, and shifted to the new temperature immediately after eclosion. 55% humidity and 12h:12h LD cycles were maintained.

Negative geotaxis was estimated in male flies as the average time required to climb a height of 9 cm inside a fly vial. Flies were house kept in vials in groups of as many as 5 flies right after eclosion. After 3-5 days, wings were manually clipped under CO2 flow, and returned to the vial for additional 48h. Then, they were transferred to an experimental vial with no food and recorded for 2-3 minutes after 3 taps. Each fly was monitored for three trials.

Fertility was measured as the proportion of flies giving rise to viable progeny. Individual males and females were crossed to 3 flies of the opposite sex in single vials. Progeny was screened in these vials one week after.

## QUANTIFICATION AND STATISTICAL ANALYSIS

Statistical significance was evaluated using Fisher’s exact test for receptivity (Figure 3, 4) and fertility (Supplementary Figure 5); Mann-Whitney non parametric test for egg-laying (Figure 3, 4), ovipositor extrusions (Figure 3, 4), vaginal plates openings (Figure 3, 4), fluorescence intensity (Figure 2, 3 and Supplementary Figure 4), number of neurons/nuclei (Figures 1, 2), number of sex combs (Supplementary Figure 5), and climbing (Supplementary Figure 5); and unpaired t-test with Welch’s correction for differential gene expression analysis (Figure 1, 2). Ns=not significant, * p<0.05, ** p<0.01, *** p<0.001, **** p<0.0001. Error bars in Figure 1, 2, 3, 4 and Supplementary Figure 5 represent SEM. All n values are displayed on the figures.

## KEY RESOURCES TABLE (KRT)

|  | SOURCE | IDENTIFIER |
| --- | --- | --- |
| **Antibodies** |  |  |
| rabbit anti-Hth (1:500) | Salvany et al, 2009 | N/A |
| guinea pig anti-Hth (1:500) | Salvany et al, 2009 | N/A |
| Mouse anti-Abd-A (1:500) | Gift of Ian Duncan | Department of<br>Biology.<br>Washington<br>University in St.<br>Louis |
| Rat anti-Dsx | Sanders and Arbeitman,<br>2008 | N/A |
| Donkey anti-rabbit Alexa fluor 488 | Thermo Fisher Scientific | A32790 |
| Goat anti-rabbit Alexa fluor 647 | Thermo Fisher Scientific | A32733 |
| Donkey anti-mouse Alexa fluor 594 | Thermo Fisher Scientific | A32744 |
| Donkey anti-mouse Alexa fluor 647 | Thermo Fisher Scientific | A32787 |
| Goat anti-guinea pig Alexa fluor 488 | Thermo Fisher Scientific | A-11073 |
| Donkey anti-rat Alexa fluor 594 | Thermo Fisher Scientific | A-21209 |
| **Chemicals, Peptides, and Recombinant Proteins** |  |  |
| Gel Loading Buffer II | Invitrogen | AM8547 |
| AccuPrime Pfx DNA Polymerase | Thermo Fisher | 12344024 |
| Trizol reagent | Life Technologies | 15596018 |
| Oligo d(T)25 Magnetic Beads | New England Biolabs | S1419S |
| Millennium™ RNA Markers | Thermo Fisher | AM7150 |
| Amersham Megaprime DNA Labeling<br>System | Cytiva | RPN1606 |
| VECTASHIELD_ Vibrance Antifade<br>Mounting Medium without DAPI | Vector Laboratories | H-1700-2 |
| VECTASHIELD_ Antifade Mounting<br>Medium with DAPI | Vector Laboratories | H-1500-10 |
| **Experimental Models: Organisms/Strains** |  |  |
| *mir[Δ]* | (Bender, 2008) Gift of<br>Welcome Bender,<br>Harvard University | N/A |
| *Canton-S* | Gift of Karla Kaun,<br>Department of<br>Neuroscience, Brown<br>University. | N/A |
| *VT-lines and SAG-1 split Gal4 line* | (Feng et al., 2014). Gift of<br>Barry Dickson, Janelia<br>Farm Research Campus. | N/A |
| *tub-GFP-sensors for miR-iab-4 and miR-iab-8* | Tyler et al., 2008, Eric Lai<br>lab | N/A |
| *hth[BSmut]* | Garaulet et al., 2020, Eric<br>Lai lab. | N/A |
| *2xUAS-dsx-RNAi[BB]* | Robinett et al., 2010 | N/A |
| *UAS-hth-RNAi* | Vienna Drosophila RNAi<br>Center | 100630 |
| *tubGal80ts* | Bloomington Drosophila<br>Stock Center | 7108 |
| *elav-Gal4[C-155]* | Bloomington Drosophila<br>Stock Center | 458 |
| *UAS-mCD8-GFP* | Bloomington Drosophila<br>Stock Center | 5137 |
| *UAS-Red-Stinger* | Bloomington Drosophila<br>Stock Center | 8547 |
| *UAS-dsx-RNAi[JF]* | Bloomington Drosophila<br>Stock Center | 26716 |
| *dsx[1]* | Bloomington Drosophila<br>Stock Center | 1679 |
| *dsx[9]* | Bloomington Drosophila<br>Stock Center | 44223 |
| *UAS-dsxF* | Bloomington Drosophila<br>Stock Center | 44223 |
| *UAS-hth* | Gift of Richard S. Mann,<br>Columbia University.<br>Casares and Mann, 1998. | N/A |
| **Deposited Data** |  |  |
| VNC RNA-seq data | This study. | NCBI GEO:<br>GSE166562 |
| Raw files of immunostaining | This study. | N/A |
| Raw files of behavior | This study. | N/A |
| **Software and algorithms:** |  |  |
| Prism 7 for Mac OS X | GraphPad | https://www.graphpad.com |
| Fiji ImageJ 2.0.0-rc-68/1.52e | N/A | https://imagej.nih.gov/ij |
| Rstudio Version 1.2.1335 | Rstudio | https://rstudio.com/ |
| IGV_2.6.3 | Broad Institute | http://software.broadinstitute.org/software/igv/ |

## Declaration of Interests

The authors declare no competing interests.

## Acknowledgments

We thank Welcome Bender, Barry Dickson, Natalia Azpiazu, Carlos Ribeiro, Carolina Rezaval, Stephen Goodwin, Karla Kaun, Michelle Arbeitman, Richard Mann, Ian Duncan and the Bloomington *Drosophila* Stock Center for fly strains, plasmids, and antibodies used in this study. We thank Ernesto Sánchez-Herrero and Paloma Martín for help and support. Work in E.C.L.’s group was supported by the NIH (R01-GM083300 and R01-NS083833) and MSK Core grant P30-CA008748.

## Author contributions

Conceptualization, D.L.G., E.C.L.; Methodology, D.L.G.; Formal analysis, D.L.G., A.M.; Investigation, D.L.G., A.M., E.C.L.; Resources, D.L.G., E.C.L.; Visualization, D.L.G.; Supervision, D.L.G., E.C.L.; Writing – original Draft and Reviewing and Editing, D.L.G., E.C.L.; Project administration, D.L.G., E.C.L.; Funding acquisition, E.C.L.

## Supplementary Figures and Table

**Supplementary Figure 1**. Transcriptome analyses of the iab-8 domain.

(A) Cartoon illustrating *mir-iab-4/8* regulation of *hth* as well as the *mir-iab-4/8* and *hth* binding sites mutants used in this study. (B) Pattern of expression of miR-iab-4 and miR-iab-8 and strategy for manual dissection of the iab-8 region of the larval ventral nerve cord (VNC), corresponding to the domain of mir-iab-8 expression. Triplicate biological samples were isolated from *Canton-S*, trans-heterozygotes of *mir-iab-4/8* deletion (*mir[Δ/C11]*), and the *hth[BSmut]* bearing mutations in all miR-iab-4/8 binding sites in its 3’ UTR. (C) MDS plot of the RNA-seq samples. (D) MA plot comparing Δmir-iab-4/8 to *Canton-S* shows relatively subtle overall gene expression changes, with little changes to direct miR-iab-8-5p targets. Amongst previously characterized direct miR-iab-8-5p targets encoding homeobox genes, both TALE cofactors (hth and exd) were substantially upregulated. (E) MA plot comparing *hth[BSmut]* to *Canton-S* also shows relatively subtle overall gene expression changes. *hth* was reproducibly upregulated, although to a lesser extent than in *Δmir-iab-4/8*, but *exd* was unchanged.

**Supplementary Figure 2**. Selected genes that are co-regulated in *Δmir-iab-4/8* and *hth[BSmut]*. Shown are example loci that are coordinately upregulated or downregulated the iab-8 region of *Δmir-iab-4/8* and *hth[BSmut]* VNC, compared to control *Canton-S*. Data are from the triplicate RNA-seq experiments.

**Supplementary Figure 3**. Highly complementary expression of Hth and Dsx in the brain. Schematic of the *Drosophila* central nervous system (Center). Double labeling for Hth (green) and Dsx (purple) in the brain (upper half) and VNC (lower half). Maximum projection of the brain is shown, revealing broad expression of Hth and sparser accumulation of Dsx. aDN, pC3 and pC1 clusters are shown in higher magnification illustrating largely complementary accumulation of these nuclear markers, even in closely apposed cells. In the VNC, very few cells present strong colocalization of both proteins. Scalebar= 40 µM.

**Supplementary Figure 4**. Characterization of SAG-1 neurons in the VNC.

(A) Images show the regulatory intersection of *VT-7068* and *VT-50405* using split-Gal4 lines, that defines a sparse pattern with typically four SAG-1 neurons within the abdominal ganglion (AbG). (B) These abdominal SAG-1 neurons project their axons to the central brain. (C) The somas of abdominal SAG-1 neurons, labeled here using nuclear Red-Stinger (red-stg, in cyan), reside in the iab-8 domain of the AbG, posterior to abd-A expressing segments (in red). (D, E) Quantification of Dsx and Hth levels in the 4 abdominal SAG-1 of control, *SAG-1>UAS-hth*, and *SAG-1>UAS-dsxF* flies. (F) Images show representative images of the quantifications shown in D and E. Mann-Whitney non-parametric test ns, not significant, *p < 0.05, ****p < 0.0001. Error bars, SEM. Scalebars in 50 μm.

**Supplementary Figure 5**. Additional characterization of *doublesex* function.

(A) Images show male sex combs on the first tarsal segment of the front leg (distal to the left, proximal to the right). Viable loss of function mutations of *dsx* result in complete loss of male features of the sex combs: thinner, less pigmented and reduced bristle numbers. Nonetheless, heterozygosity for *dsx* does not decrease male sex combs number or appearance. A published double *dsx-RNAi* transgene from Bruce Baker’s lab (“BB”) does not affect sex comb formation at 25°C, as reported, but expression of an RNAi transgene from the TRiP-JF collection mildly reduces their numbers. (B) Quantification of male sex comb numbers in controls, mutants, and knockdown genotypes, all at 25°C. (C, D) Behavioral analysis of locomotor ability and fertility, showing that Dsx levels do not affect these behaviors. (E) Images show abdominal SAG-1 lineages in wild-type controls (*Canton-S*) and the two *dsx* knockdown conditions. No gross anatomy defects were found in the number, location or dendrite arborization at the VNC nor in their axonal pattern in the brain. Mann-Whitney non-parametric test (B,C) Fisher’s exact test (D). ns, not significant, ****p < 0.0001. Error bars, SEM. Scalebar 50 μm.

**Supplementary Table 1**. Differential gene expression analysis in iab-8 region of *Δmir* and *hth[BSmut]* VNC.

## Notes

### Competing Interest Statement

The authors have declared no competing interest.

### Summary of Updates

New immunostaining data was added to Figures 1-3. New behavioral analyses were added to Figure 3-4. New supplemental figures were added.

## References

Agarwal, V., Subtelny, A.O., Thiru, P., Ulitsky, I., and Bartel, D.P. (2018). Predicting microRNA targeting efficacy in Drosophila. Genome biology 19, 152.

Anholt, R.R.H., O’Grady, P., Wolfner, M.F., and Harbison, S.T. (2020). Evolution of Reproductive Behavior. Genetics 214, 49–73.

Bender, W. (2008). MicroRNAs in the Drosophila bithorax complex. Genes & development 22, 14–19.

Durinck, S., Moreau, Y., Kasprzyk, A., Davis, S., De Moor, B., Brazma, A., and Huber, W. (2005). BioMart and Bioconductor: a powerful link between biological databases and microarray data analysis. Bioinformatics 21, 3439–3440.

Feng, K., Palfreyman, M.T., Hasemeyer, M., Talsma, A., and Dickson, B.J. (2014). Ascending SAG neurons control sexual receptivity of Drosophila females. Neuron 83, 135–148.

Garaulet, D.L., Castellanos, M.C., Bejarano, F., Sanfilippo, P., Tyler, D.M., Allan, D.W., Sanchez-Herrero, E., and Lai, E.C. (2014). Homeotic Function of Drosophila Bithorax-Complex miRNAs Mediates Fertility by Restricting Multiple Hox Genes and TALE Cofactors in the CNS. Developmental cell 29, 635–648.

Garaulet, D.L., Zhang, B., Wei, L., Li, E., and Lai, E.C. (2020). miRNAs and Neural Alternative Polyadenylation Specify the Virgin Behavioral State. Developmental cell 54, 410–423.

Gummalla, M., Maeda, R.K., Castro Alvarez, J.J., Gyurkovics, H., Singari, S., Edwards, K.A., Karch, F., and Bender, W. (2012). abd-A Regulation by the iab-8 Noncoding RNA. PLoS genetics 8, e1002720.

Hasemeyer, M., Yapici, N., Heberlein, U., and Dickson, B.J. (2009). Sensory neurons in the Drosophila genital tract regulate female reproductive behavior. Neuron 61, 511–518.

Jang, Y.H., Chae, H.S., and Kim, Y.J. (2017). Female-specific myoinhibitory peptide neurons regulate mating receptivity in Drosophila melanogaster. Nature communications 8, 1630.

Kim, D., Langmead, B., and Salzberg, S.L. (2015). HISAT: a fast spliced aligner with low memory requirements. Nature methods 12, 357–360.

Kopp, A. (2012). Dmrt genes in the development and evolution of sexual dimorphism. Trends in genetics : TIG 28, 175–184.

Kubli, E., and Bopp, D. (2012). Sexual behavior: how Sex Peptide flips the postmating switch of female flies. Curr Biol 22, R520–522.

Liao, Y., Smyth, G.K., and Shi, W. (2019). The R package Rsubread is easier, faster, cheaper and better for alignment and quantification of RNA sequencing reads. Nucleic acids research 47, e47.

McRobert, S.P., and Tompkins, L. (1985). The effect of transformer, doublesex and intersex mutations on the sexual behavior of Drosophila melanogaster. Genetics 111, 89–96.

Mezzera, C., Brotas, M., Gaspar, M., Pavlou, H.J., Goodwin, S.F., and Vasconcelos, M.L. (2020). Ovipositor Extrusion Promotes the Transition from Courtship to Copulation and Signals Female Acceptance in Drosophila melanogaster. Curr Biol 30, 3736–3748 e3735.

Nothiger, R., Leuthold, M., Andersen, N., Gerschwiler, P., Gruter, A., Keller, W., Leist, C., Roost, M., and Schmid, H. (2009). Genetic and developmental analysis of the sex-determining gene ‘double sex’ (dsx) of Drosophila melanogaster. Genetics Research 50, 113–123.

Ogawa, S., and Makino, J. (1984). Aggressive behavior in inbred strains of mice during pregnancy. Behavioral and neural biology 40, 195–204.

Pai, C.Y., Kuo, T.S., Jaw, T.J., Kurant, E., Chen, C.T., Bessarab, D.A., Salzberg, A., and Sun, Y.H. (1998). The Homothorax homeoprotein activates the nuclear localization of another homeoprotein, extradenticle, and suppresses eye development in Drosophila. Genes & development 12, 435–446.

Rezaval, C., Pavlou, H.J., Dornan, A.J., Chan, Y.B., Kravitz, E.A., and Goodwin, S.F. (2012). Neural circuitry underlying Drosophila female postmating behavioral responses. Curr Biol 22, 1155–1165.

Rideout, E.J., Dornan, A.J., Neville, M.C., Eadie, S., and Goodwin, S.F. (2010). Control of sexual differentiation and behavior by the doublesex gene in Drosophila melanogaster. Nature neuroscience 13, 458–466.

Rieckhof, G.E., Casares, F., Ryoo, H.D., Abu-Shaar, M., and Mann, R.S. (1997). Nuclear translocation of extradenticle requires homothorax, which encodes an extradenticle-related homeodomain protein. Cell 91, 171–183.

Robinett, C.C., Vaughan, A.G., Knapp, J.M., and Baker, B.S. (2010). Sex and the single cell. II. There is a time and place for sex. PLoS biology 8, e1000365.

Robinson, M.D., McCarthy, D.J., and Smyth, G.K. (2010). edgeR: a Bioconductor package for differential expression analysis of digital gene expression data. Bioinformatics 26, 139–140.

Salvany, L., Aldaz, S., Corsetti, E., and Azpiazu, N. (2009). A new role for hth in the early pre- blastodermic divisions in drosophila. Cell cycle 8, 2748–2755.

Sanders, L.E., and Arbeitman, M.N. (2008). Doublesex establishes sexual dimorphism in the Drosophila central nervous system in an isoform-dependent manner by directing cell number. Developmental biology 320, 378–390.

Schindelin, J., Arganda-Carreras, I., Frise, E., Kaynig, V., Longair, M., Pietzsch, T., Preibisch, S., Rueden, C., Saalfeld, S., Schmid, B., et al. (2012). Fiji: an open-source platform for biological-image analysis. Nature methods 9, 676–682.

Soller, M., Haussmann, I.U., Hollmann, M., Choffat, Y., White, K., Kubli, E., and Schafer, M.A. (2006). Sex-peptide-regulated female sexual behavior requires a subset of ascending ventral nerve cord neurons. Curr Biol 16, 1771–1782.

Svare, B., Mann, M.A., Broida, J., and Michael, S.D. (1982). Maternal aggression exhibited by hypophysectomized parturient mice. Horm Behav 16, 455–461.

Tyler, D.M., Okamura, K., Chung, W.J., Hagen, J.W., Berezikov, E., Hannon, G.J., and Lai, E.C. (2008). Functionally distinct regulatory RNAs generated by bidirectional transcription and processing of microRNA loci. Genes & development 22, 26–36.

Wang, F., Wang, K., Forknall, N., Parekh, R., and Dickson, B.J. (2020a). Circuit and Behavioral Mechanisms of Sexual Rejection by Drosophila Females. Curr Biol 30, 3749–3760 e3743.

Wang, F., Wang, K., Forknall, N., Patrick, C., Yang, T., Parekh, R., Bock, D., and Dickson, B.J. (2020b). Neural circuitry linking mating and egg laying in Drosophila females. Nature 579, 101–105.

Wang, K., Wang, F., Forknall, N., Yang, T., Patrick, C., Parekh, R., and Dickson, B.J. (2021). Neural circuit mechanisms of sexual receptivity in Drosophila females. Nature 589, 577–581.

Yang, C.H., Rumpf, S., Xiang, Y., Gordon, M.D., Song, W., Jan, L.Y., and Jan, Y.N. (2009). Control of the postmating behavioral switch in Drosophila females by internal sensory neurons. Neuron 61, 519–526.

