## Supplemental Figures 1-5 for "A double negative post-transcriptional regulatory circuit underlies the virgin behavioral state"

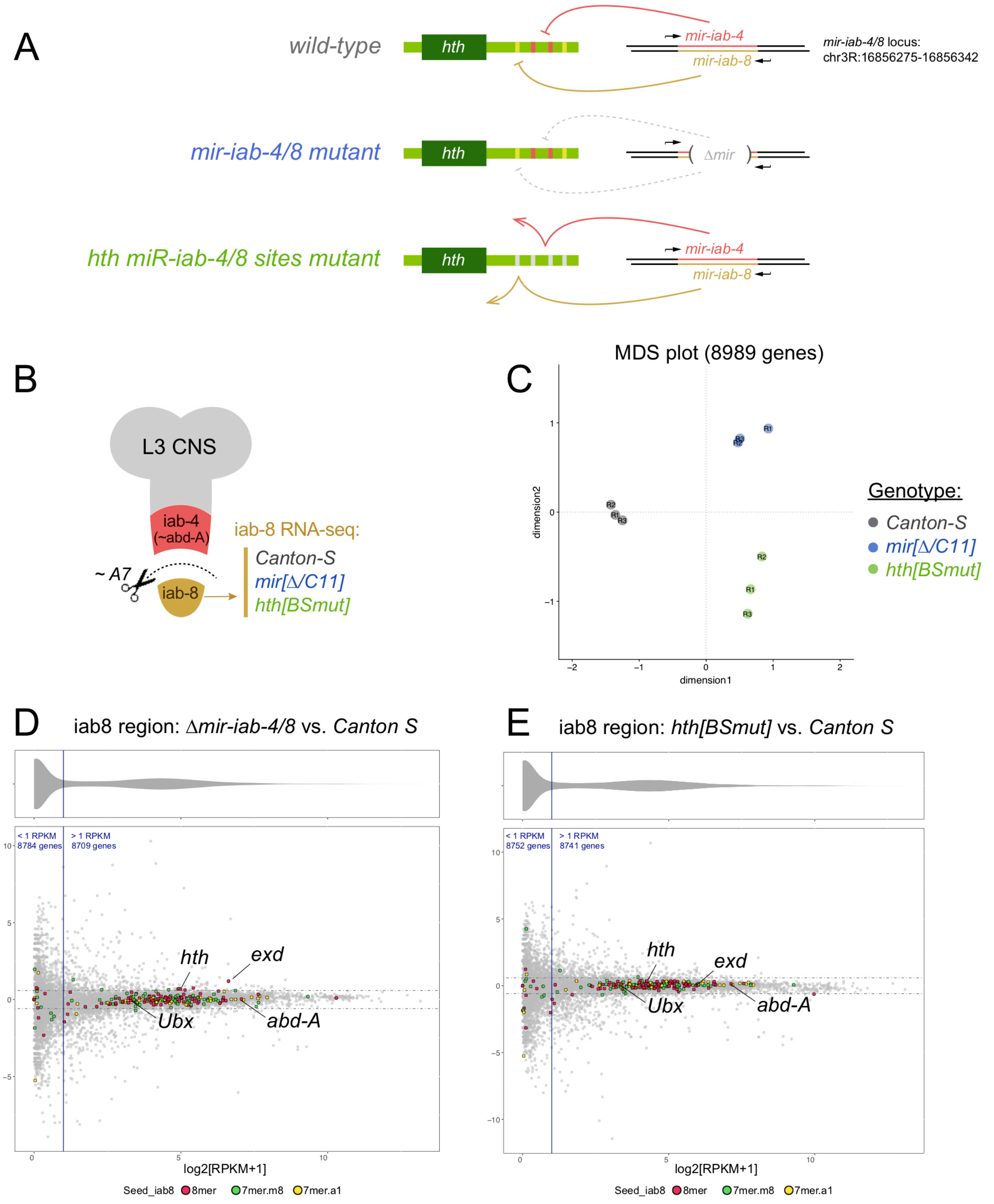


Supplementary Figure 1. Transcriptome analyses of the iab-8 domain.

(A) Cartoon illustrating *mir-iab-4/8* regulation of hth as well as the *mir-iab-4/8* and *hth* binding sites mutants used in this study. (B) Pattern of expression of miR-iab-4 and miR-iab-8 and strategy for manual dissection of the iab-8 region of the larval ventral nerve cord (VNC), corresponding to the domain of *mir-iab-8* expression. Triplicate biological samples were isolated from *Canton-S*, trans-heterozygotes of *mir-iab-4/8* deletion (*mir[Δ/C11]*), and the *hth[BSmut]* bearing mutations in all miR-iab-4/8 binding sites in its 3' UTR. (C) MDS plot of the RNA-seq samples. (D) MA plot comparing Δ*mir-iab-4/8* to *Canton-S* shows relatively subtle overall gene expression changes, with little changes to direct miR-iab-8-5p targets. Amongst previously characterized direct miR-iab-8-5p targets encoding homeobox genes, both TALE cofactors (*hth* and *exd*) were substantially upregulated. (E) MA plot comparing *hth[BSmut]* to *Canton-S* also shows relatively subtle overall gene expression changes. *hth* was reproducibly upregulated, although to a lesser extent than in Δ*mir-iab-4/8*, but *exd* was unchanged.

Garaulet et al
Supplementary Figure 1

### Example genes upregulated in both Δ*mir-iab-4/8* and *hth[BSmut]*

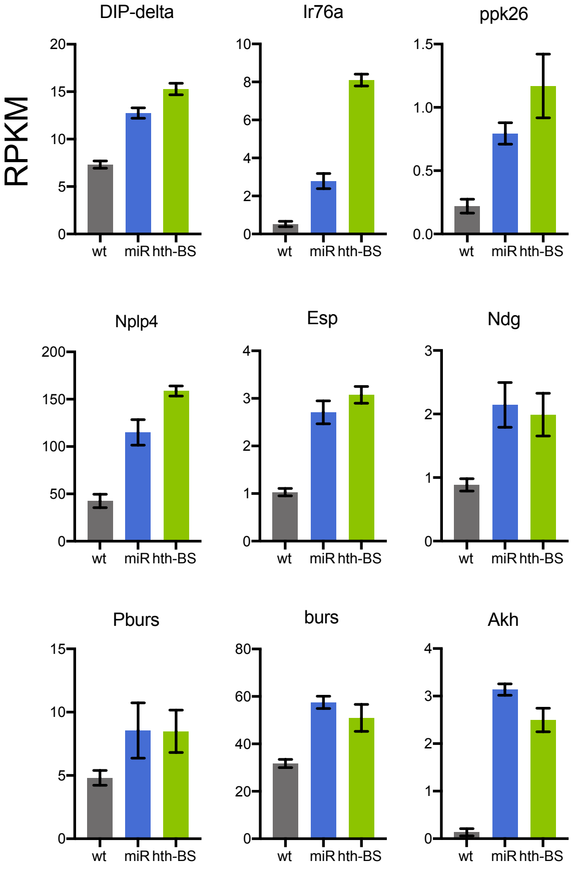


| | |
|---|---|
| DIP-delta | Neural |
| Ir76a | Neural. Ionotropic receptor. Expressed in antennal sensory neurons |
| ppk26 | Sodium channel family . Ppk neurons involved in switch |
| Nplp4 | Neuropeptide like percursors, neuropeptide |
| Esp | Involved in female remating receptivity behavior |
| Ndg | Neural. Basal membrane and ECM. Involved in synaptic plasticity |
| Pburs | Partner of Burst - Neural exclusive |
| Burs | Neurohormone - not very neuron specific |
| Akh | Adipokinetic hormone. High neural |

### Example genes downregulated in both Δ*mir-iab-4/8* and *hth[BSmut]*

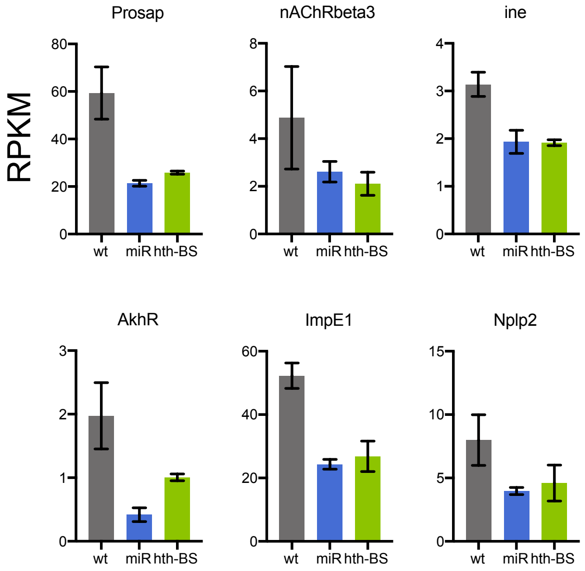


| | |
|---|---|
| Prosap | High in neurons. Synaptic growth, NMJ |
| nAChRbeta3 | nicotinic Acetylcholine Receptor |
| ine | High in CNS (mostly adults). SLC6A family of neurotransmitter transporters |
| AkhR | protein-coupled receptor for the hormone encoded by Akh |
| Flo2 | CNS-enriched scaffold protein, involved in wg and hh spreading |
| ImpE1 | Ecdysone-inducible gene E1. CNS disc exclusive |
| Nplp2 | Neuropeptide-like precursor 2. Neurohormone |

Supplementary Figure 2. Selected genes that are co-regulated in Δ*mir-iab-4/8* and *hth[BSmut]*.

Shown are example loci that are coordinately upregulated or downregulated the iab-8 region of Δ*mir-iab-4/8* and *hth[BSmut]* VNC, compared to control *Canton-S*. Data are from the triplicate RNA-seq experiments.

Garaulet et al
Supplementary Figure 2

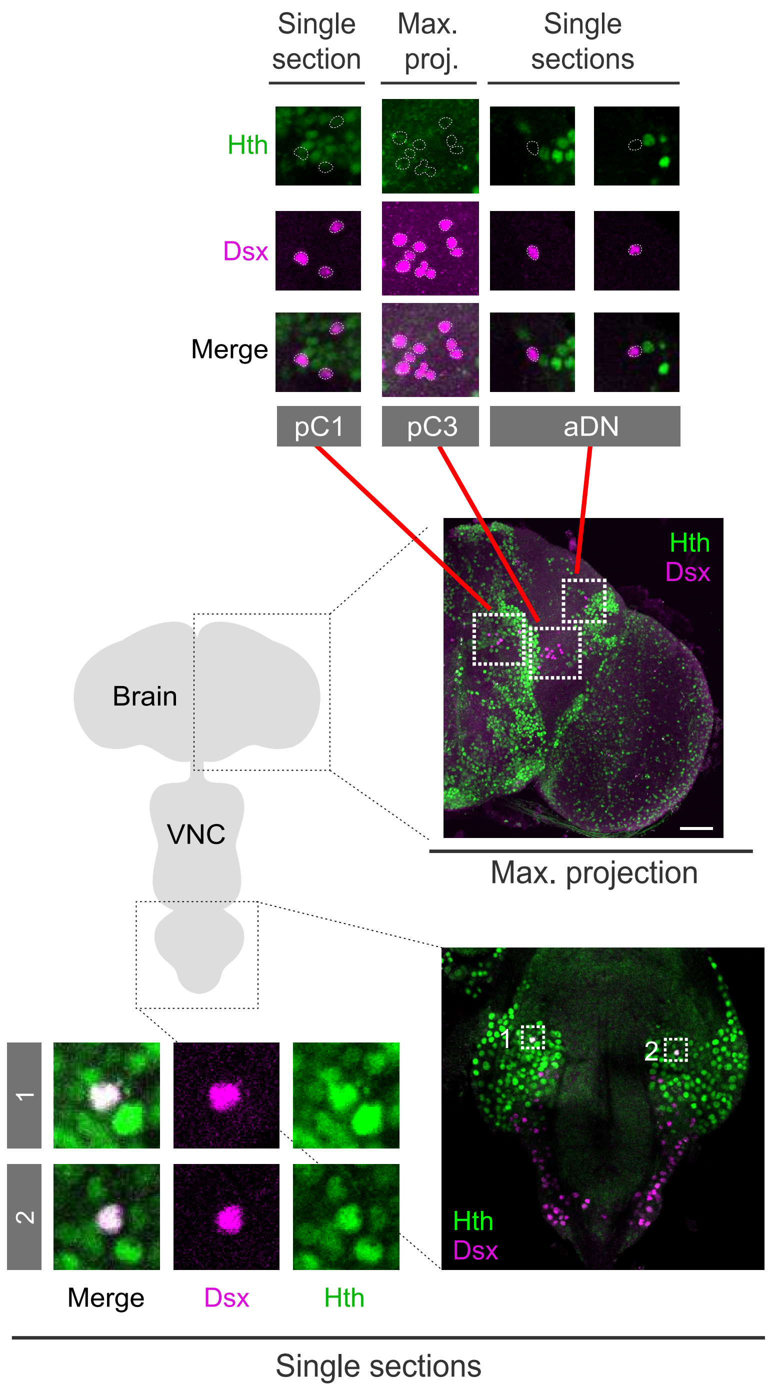


Supplementary Figure 3. Highly complementary expression of Hth and Dsx in the brain.

Schematic of the *Drosophila* central nervous system (Center). Double labeling for Hth (green) and Dsx (purple) in the brain (upper half) and VNC (lower half). Maximum projection of the brain is shown, revealing broad expression of hth and sparser accumulation of dsx. aDN, pC3 and pC1 clusters are shown in higher magnification illustrating largely complementary accumulation of these nuclear markers, even in closely apposed cells. In the VNC, very few cells present strong colocalization of both proteins. Scalebar= 40uM.

Garaulet et al
Supplementary Figure 3

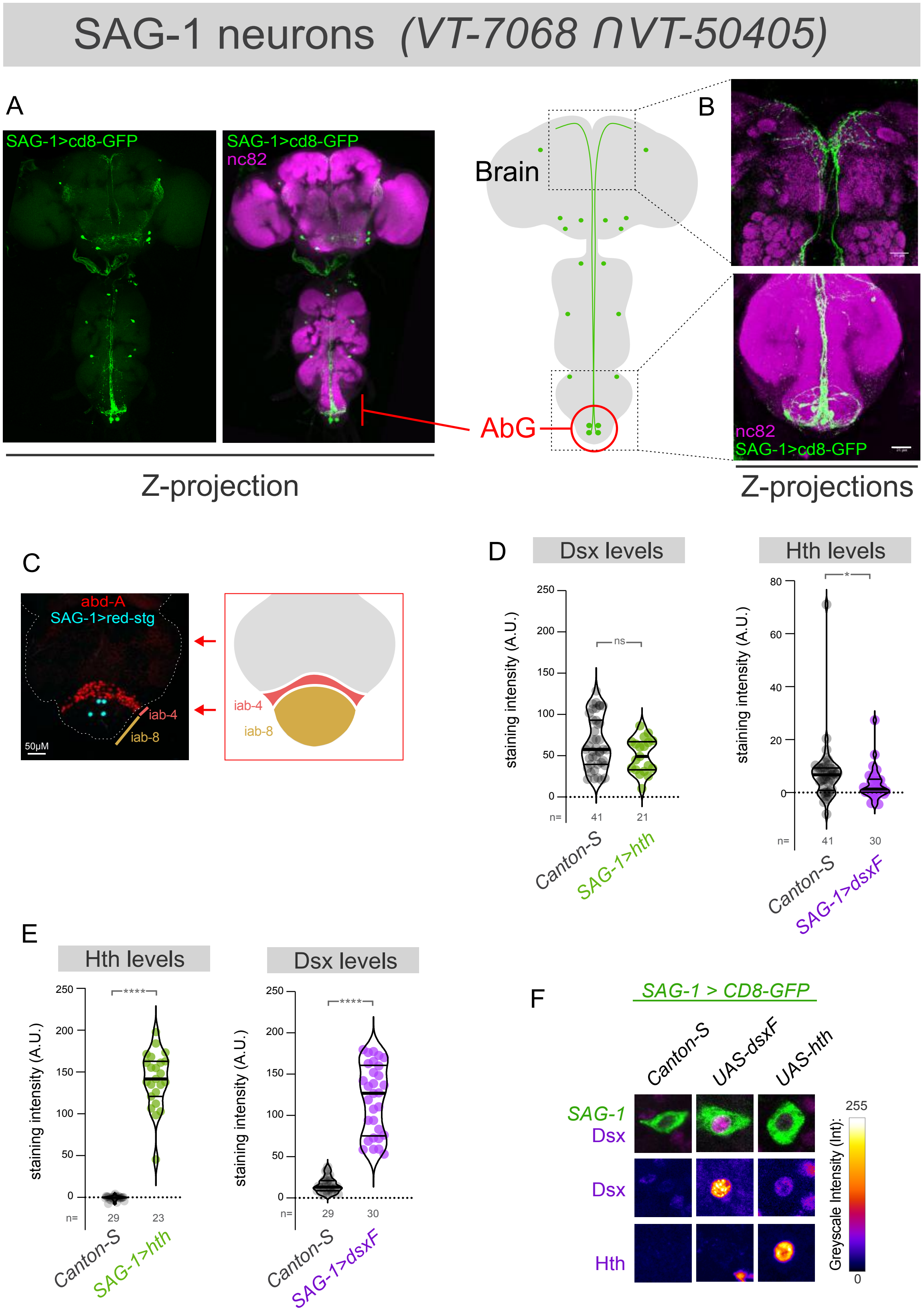


Supplementary Figure 4. Characterization of SAG-1 neurons in the VNC.

(A) Images show the regulatory intersection of *VT-7068* and *VT-50405* using split-Gal4 lines, that defines a sparse pattern with typically four SAG-1 neurons within the abdominal ganglion (AbG). (B) These abdominal SAG-1 neurons project their axons to the central brain. (C) The somas of abdominal SAG-1 neurons, labeled here using nuclear Red-Stinger (red-stg, in cyan), reside in the iab-8 domain of the AbG, posterior to abd-A expressing segments (in red). (D, E) Quantification of Dsx and Hth levels in the 4 abdominal SAG-1 of control flies, *SAG-1>UAS-hth*, and *SAG-1>UAS-dsxF* flies. (F) Images show representative images of the quantifications shown in D and E. Mann-Whitney non parametric test ns, not significant, *$p < 0.05$, ****$p < 0.0001$. Error bars, SEM. Scalebars in 50 μm.

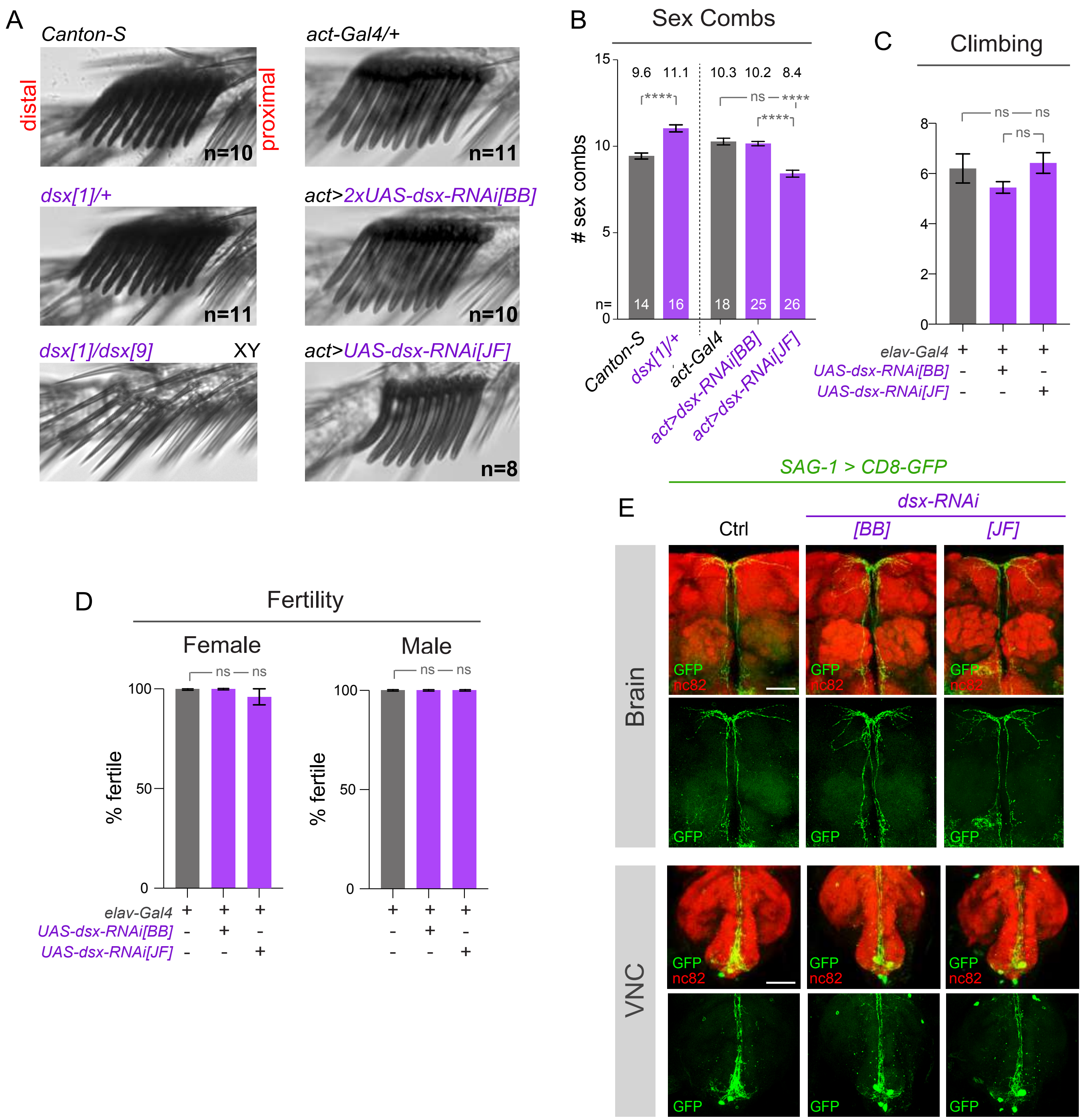


Supplementary Figure 5. Additional characterization of *doublesex* function.

(A) Images show male sex combs on the first tarsal segment of the front leg (distal to the left, proximal to the right). Viable loss of function mutations of *dsx* result in complete loss of male features of the sex combs: thinner, less pigmented and reduced bristle numbers. Nonetheless, heterozigosity for *dsx* does not decrease male sex combs number or appearance. A published double *dsx-RNAi* transgene from Bruce Baker's lab ("BB") does not affect sex comb formation at 25ºC, as reported, but expression of an RNAi transgene from the TRiP-JF collection mildly reduces their numbers. (B) Quantification of male sex comb numbers in controls, mutants, and knockdown genotypes, all at 25ºC. (C, D) Behavioral analysis of locomotor ability and fertility, showing that Dsx levels do not affect these behaviors. (E) Images show abdominal SAG-1 lineages in wild-type controls (*Canton-S*) and the two *dsx* knockdown conditions. No gross anatomy defects were found in the number, location or dendrite arborization at the VNC nor in their axonal pattern in the brain. Mann-Whitney non parametric test (B,C) Fisher's exact test (D). ns, not significant, ****$p < 0.0001$. Error bars, SEM. Scalebar 50 µm.

Garaulet et al
Supplementary Figure 5
